# Building functional circuits in multispecies brains

**DOI:** 10.1101/2023.04.13.536815

**Authors:** Benjamin T. Throesch, Muhammad Khadeesh bin Imtiaz, Rodrigo Muñoz-Castañeda, Masahiro Sakurai, Andrea L. Hartzell, Kiely N. James, Alberto R. Rodriguez, Greg Martin, Giordano Lippi, Sergey Kupriyanov, Zhuhao Wu, Pavel Osten, Juan Carlos Izpisua Belmonte, Jun Wu, Kristin K. Baldwin

## Abstract

The genome is the ultimate architect of the brain. Its evolutionary variations build the neural circuits that endow each species with its innate senses and behaviors. A central question for neuroscience and translational medicine is whether neural circuits from two species can be made to function in an intact brain. Here, we establish genetic tools and use blastocyst complementation to selectively build and test interspecies neural circuits in rat-mouse brains. Despite ∼10-20 million years of evolution and prominent differences in brain size and cellular composition, rat pluripotent stem cells injected into mouse blastocysts widely populate and persist in the mouse brain. Unexpectedly, the mouse niche reprograms the birthdates of cortical and hippocampal rat neurons, where they also form synaptically active rat-mouse circuits. By genetically disabling host olfactory circuitry, we show that rat neurons restore synaptic information flow from the nose to the cortex. Rat neurons can also rescue a primal olfactory behavior (food-seeking), though less than mouse controls. By enabling a mouse to sense the world with rat neurons, we highlight the power of interspecies neural blastocyst complementation to uncover mechanisms of neural circuit development and evolution, and to inform efforts to rescue neural circuits affected by injury or disease.

---

Evolution has sculpted brain development to enable the precise assembly of diverse cellular subtypes into neural circuits with species-specific functions. Interspecies chimeras generated from either induced pluripotent stem cells (iPSCs) or embryonic stem cells (ESCs) hold tremendous potential to systematically dissect processes underlying neuronal development and circuit formation across species from the earliest stages in an intact, injury-free whole organ system. Interspecies chimeras between several rodent species have been reported with donor PSCs exhibiting broad but apparently random contribution to different tissues throughout the host organism^1–5^. To enrich donor PSCs in a specific tissue or organ within or across different species, a method called blastocyst complementation has been developed^6–10^ in which the host animal’s developmental niche is emptied using gene knockouts or genetic cell ablation. Recently this approach has been applied to study forebrain development in mouse-mouse chimeras^11^. This intraspecies neural blastocyst complementation study showed that ablation of cortical neurons through selective expression of diphtheria toxin subunit A (DTA) enabled donor cells to populate the vacant developmental niche in the host embryo, thereby producing mouse chimeras with brains built from two different genetic backgrounds. While similar cross species transplantation experiments have been widely informative in studying brain development, none of these have interrogated the capacity for cells of two species to interact in the blastocyst and progress all the way to forming specific neural circuits in the brain.

To date, neural blastocyst complementation has not been studied in an interspecies setting, and when and where species-specific differences pose barriers to functional reconstitution of different neural circuits has not been addressed. To fill the gap, here, we establish interspecies brain chimeras produced through blastocyst complementation. We demonstrate that this constitutes a versatile and robust platform to explore circuit formation in an evolutionary and developmental context. Furthermore, we demonstrate the utility of different genetic strategies to evaluate the capacities and limitations of interspecies neural blastocyst complementation.

## Quantitative whole-brain imaging of rat-mouse chimeras

To quantify which regions of the mouse brain can support rat neuronal development, we generated interspecies chimeras using a rat ESC line (DAC2)^12^ that expresses a bright red-orange fluorescent protein, Kusabira Orange (KsO)^3^. We injected these ESCs into mouse blastocysts, which were then transferred into the uteri of surrogate mouse mothers to complete their development (Fig. 1a, Table 1). Rat-mouse chimeras exhibited varying degrees of rat cell contribution based on KsO fluorescence, reflecting the expected stochastic nature of chimera experiments (Fig. 1a). Chimeras with brain contribution showed non-symmetric, bilateral patterning of neural cells across brain regions including the olfactory bulbs (OBs), piriform cortex (PCx), multiple neocortical areas, hippocampus (Hipp), caudoputamen (CP), and cerebellum (CB) (Fig. 1b). Within each region, rat neurons displayed diverse yet regionally characteristic cellular morphologies and layering (Fig. 1b). Although we detected clear rat contribution throughout the chimeric brain, in some regions the KsO signal was also found in non-neuronal cell types such as microglia, glia, pericytes, and vascular endothelial cells.

**Figure 1:**
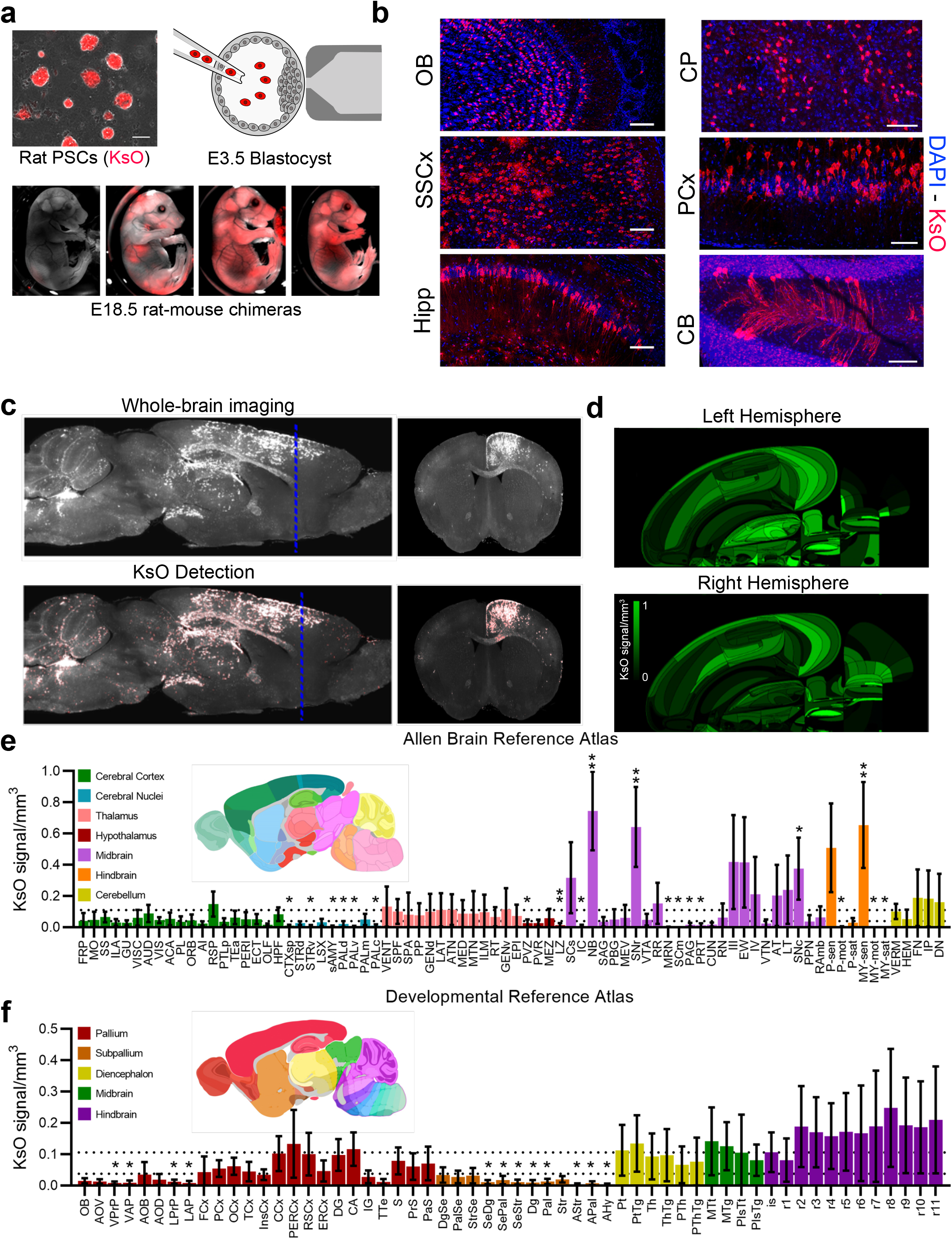
Rat cells contribute widely to the mouse brain. **a,** Schematic overview of rat-mouse chimera formation using rat pluripotent stem cells (PSCs). Rat PSCs fluorescently labelled with KsO were injected into E3.5 mouse blastocysts. Derived E18.5 fetuses displayed variable degrees of KsO contribution (shown in red), scale bar 100 μm. **b,** Representative images depict specialized, non-symmetrical contribution of rat cells to diverse neural circuits. Shown are the nuclei (DAPI, blue) and rat KsO cells (red). OB, olfactory bulb; PCx, piriform cortex; SSCx, somatosensory cortex; CP, caudoputamen; Hipp, hippocampus; CB, cerebellum. Scale bar, 100 μm. **c,** Whole-brain imaging of high-contribution chimeric brains. Representative images show KsO expression (red) detection. **d,** Flat map projections onto a registered atlas. n = 6 animals. **e,** Volumetric analysis of KsO expression in brain regions defined by the Allen Mouse Brain Reference Atlas. **f,** Volumetric analysis based on the Allen Developing Mouse Brain Reference Atlas. Results of 12 hemispheres from 6 independent brain samples are shown; mean ± 95% CI. Dashed lines indicate the 95% CI for the mean hemisphere signal. Significance was tested by repeated measures One-Way ANOVA with Dunnett’s multiple comparisons test to the mean hemisphere signal, \**p* < 0.05, \*\**p* < 0.01.

**Table 1.**
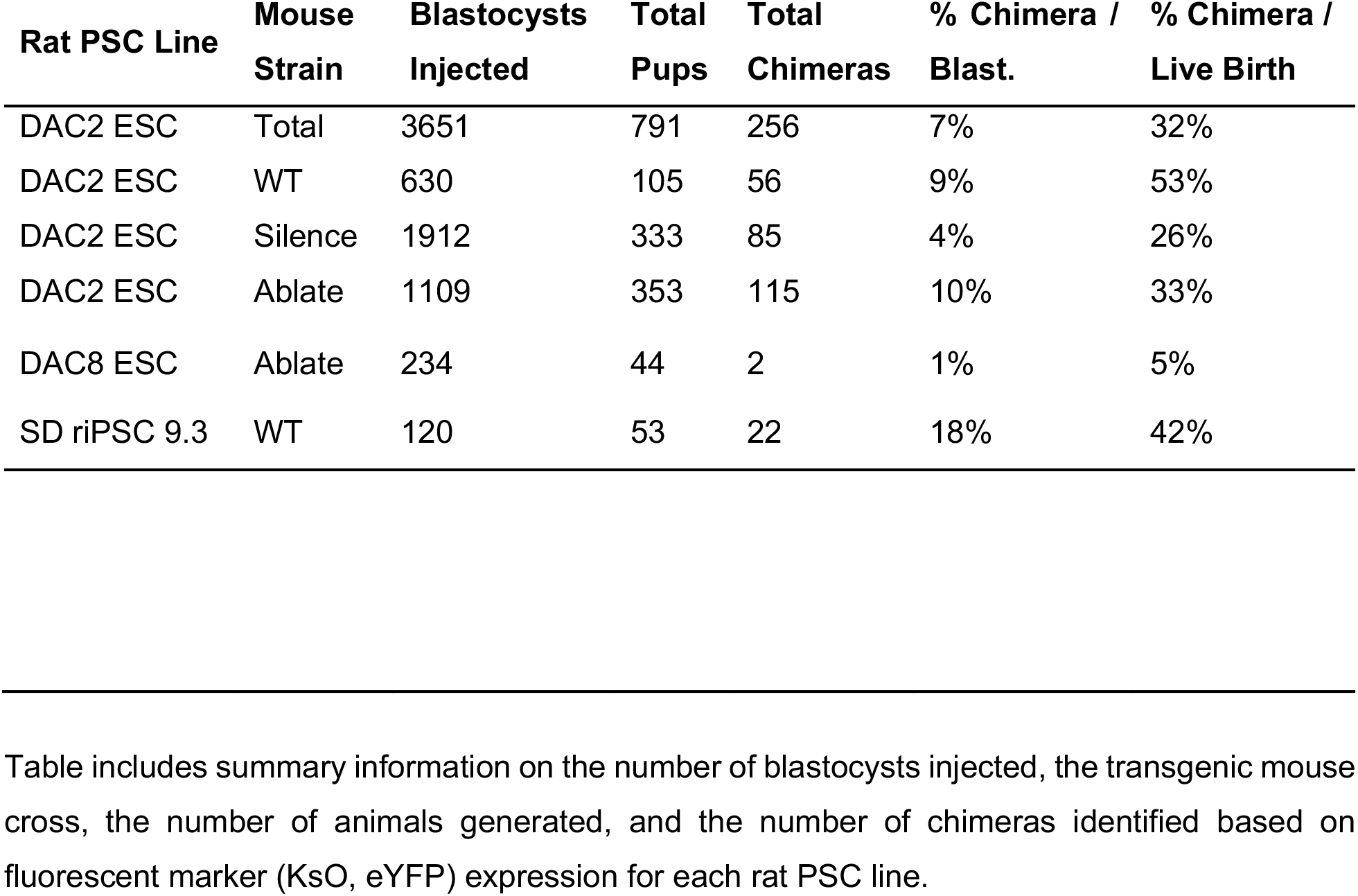
Blastocyst Complementation Efficiencies.

We performed whole brain clearing on six rat ESC-derived chimeras with high brain contribution and analyzed the extent and location of rat cells across the entire brain using light sheet microscopy (Fig. 1c, Supplementary Fig. 1a-c, Supplementary Video 1, 2, 3). To accurately quantify the contribution of rat cells across defined brain regions we developed a method for registration and quantification of the percent KsO signal per volume for any defined three-dimensional brain region. First, we analyzed average and individual chimeric brain contributions in a flat map representation (Fig. 1d, Supplementary Fig. 1c). These analyses show that rat brain contribution is broad yet highly variable among chimeras and between two hemispheres of the same animal (Supplementary Fig. 1c, d). To further quantify and identify regions of potential rat exclusion or overrepresentation, we examined two sets of brain regions as defined by the Allen Mouse Brain Reference Atlas (Fig. 1e) and the Developing Mouse Brain Reference Atlas (Fig. 1f) and plotted the average contribution across both hemispheres for the six imaged chimeric brains. Strikingly, very few regions (31/838, 3.70%) from the Brain Reference Atlas entirely lacked rat contribution. The majority of the brain regions had detectable rat signal (KsO) ranging from 0.01% to 87.12%, (Fig. 1d, Supplementary Table 1). Areas within the midbrain and hindbrain showed greater than average rat cell densities whereas cerebral nuclei and hypothalamic regions exhibited less (Fig. 1e). We also aligned the whole-brain reconstructions to the Allen Developing Mouse Brain Reference Atlas (Fig. 1f). Multiple areas of subpallium (and some pallial areas) had significantly reduced rat contribution, whereas diencephalon, midbrain, and hindbrain showed trends towards greater rat cell contribution. Together these data predict that rat cells can contribute to the vast majority of neural circuits in mouse brains. We believe this is a strong argument for developing more refined methods of interrogating cell function in interspecies brains, such as those we establish in this study.

## Reprogramming of rat neuronal birthdates

A main evolutionary distinction between brains of different species is the timing of specification and differentiation of cognate neurons. For example, rat brains are larger than mouse brains, with some expanded cortical layers, and cognate cortical neurons develop roughly one-two days later than the mouse^13^. Functional neuronal integration and circuit reconstitution might be improved if xenogeneic cells adapt to the developmental timing of the host species. However, decades of cross-species tissue grafting experiments have shown that in most contexts, grafted cells maintain their own species-specific developmental timing despite maturing in an accelerated host environment^14–17^. However, more recent heterochronic intraspecies transplantation studies have shown that some mouse cortical progenitor populations retain the capacity to read the host mouse cues and adapt to their environments^18^. Here, we sought to determine whether rat neurons developing in tandem with mouse cells from the earliest stages of embryogenesis would successfully interpret and integrate extracellular timing cues resulting in host based “reprogramming” of donor birthdates.

To examine these possibilities, we birth dated rat neurons using BrdU and EdU injections at several timepoints (ranging from E12.5 to E17.5) where birthdates of mouse and rat cells are known to differ (Fig. 2a). First, we examined the layering of rat and mouse neurons born at the same time in the cortex. Cortical regions comprise multiple layers that are built by radial migration from the ventricular zone such that early-born neurons are located in deep layers and later-born neurons migrate further to build superficial layers^19,20^. If rat neurons maintain their species intrinsic clock, EdU labeling should show a difference in the layering of mouse and rat neurons in the same chimeric brain (Fig. 2a). Alternatively, if donor (rat) neurogenesis is reprogrammed by the host mouse developmental niche, rat neurons should be intermingled spatially with mouse neurons born on the same day (Fig. 2a).

**Figure 2:**
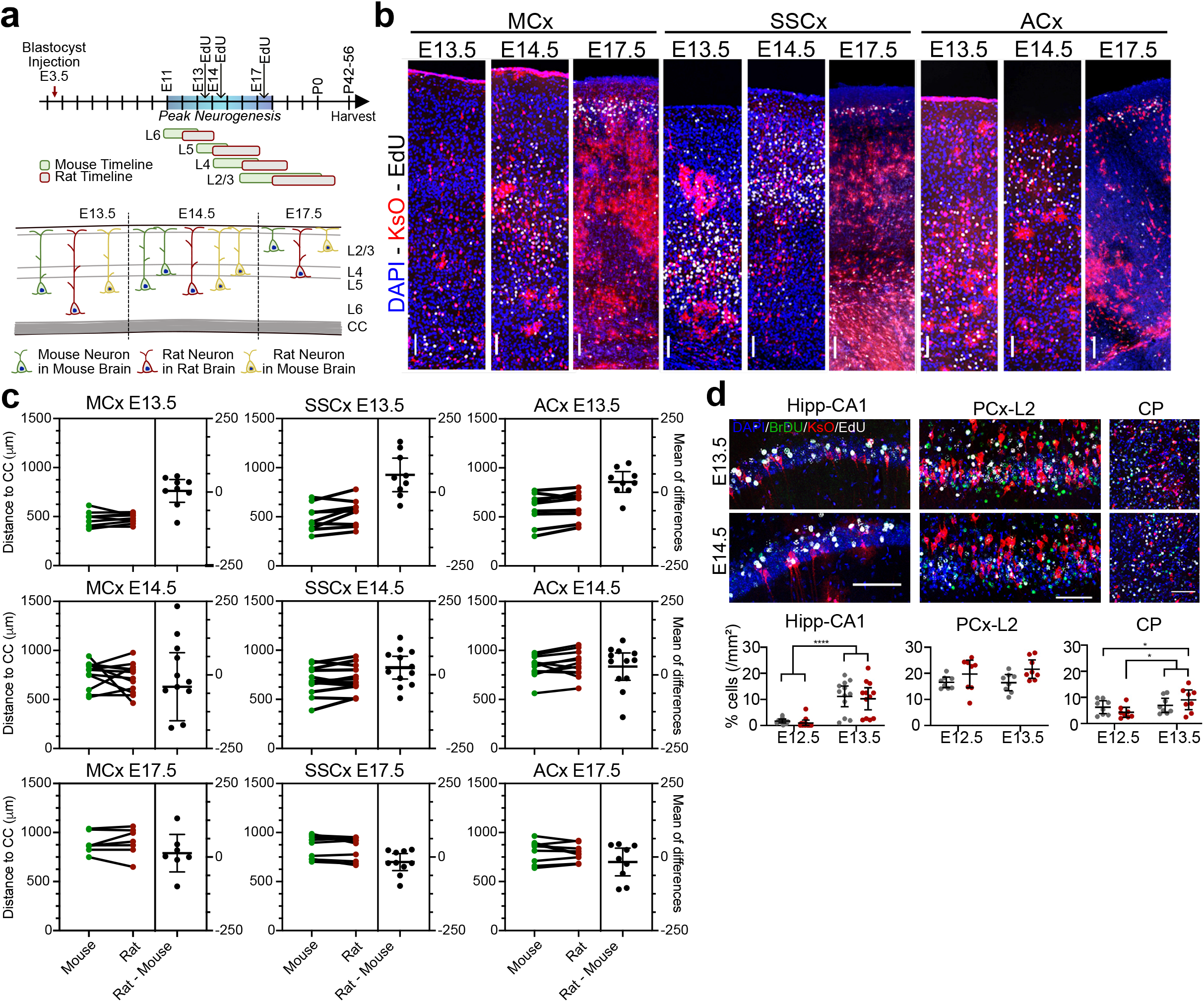
Development of rat cells within mouse neural circuits. **a,** Schematic of birth dating experiments. Surrogate mothers were injected with BrdU at E12.5 and EdU at E13.5, E14.5 or E17.5 to label different populations of neurons in chimeric embryos during peak mouse neurogenesis. Corticogenesis in rat occurs later than in mice, as in the timeline. Rat neurons could either maintain their developmental timeline (red neurons) or reprogram their timeline to that of the mouse cells (green), which is depicted in yellow (meaning rat changed to mouse timeline). **b,** Representative images of EdU labelling at E13.5, E14.5 and E17.5 in 3 different cortical regions, namely motor cortex (MCx), somatosensory cortex (SSCx) and auditory cortex (ACx). Shown are nuclei (DAPI, blue), rat KsO cells (red) and EdU labelled cells (white). Scale bar 100 μm. **c,** Rat developmental time matches that of the mouse in each region and timepoint as quantified by measuring the distance of mouse (green) and rat (red) EdU cells from the border between cortex and corpus callosum (CC). Estimation plots are shown where the measured distances for the rat and mouse cells within the same section were averaged and then paired (left). Each dot represents a section. Each plot also includes the difference between the average distance for rat and mouse cells per slice (right). T-tests on each region and timepoint were carried out using and corrected for by Bonferroni-Dunn method where n = 3, since these did not reach significance, we did not perform the more stringent test of n = 9. The number of animals per timepoint is as follows: E13.5 = 3, E14.5 = 4, E17.5 = 1. Number of sections analyzed are as follows: ACx E13.5 = 9, ACx E14.5 = 12, ACx E17.5 = 9, MCx E13.5 = 9, MCx E14.5 = 12, MCx E17.5 = 7, SSCx E13.5 = 9, SSCx E14.5 = 12, SSCx E17.5 = 10. The unadjusted p-values are as follows: ACx E13.5 = 0.05, ACx E14.5 = 0.20, ACx E17.5 = 0.43, MCx E13.5 = 0.82, MCx E14.5 = 0.46, MCx E17.5 = 0.64, SSCx E13.5 = 0.04, SSCx E14.5 = 0.19, MCx E17.5 = 0.22. The corrected p-values are as follows: ACx E13.5 = 0.15, ACx E14.5 = 0.62, ACx E17.5 > 0.99, MCx E13.5 > 0.99, MCx E14.5 > 0.99, MCx E17.5 > 0.99, SSCx E13.5 = 0.13, SSCx E14.5 = 0.58, MCx E17.5 = 0.65. **d,** Rat temporal development is largely reprogrammed in mouse non-cortical regions. Top: representative images of BrdU (labelled E12.5) and EdU (labelled E13.5 or E14.5) expression in different non-cortical regions of chimeric brains; CA1 hippocampus (Hipp-CA1), layer 2 piriform cortex (PCx-L2), caudoputamen (CP). Shown are the nuclei (DAPI, blue), BrdU positive cells (green), rat KsO cells (red) and EdU (white). Scale bar, 100 μm. Bottom: Data shown are mean ± 95% CI, n = 3-4 animals, 3 slices/animal. Significance was tested by Two-way ANOVA followed by Sidak’s multiple comparisons test, \**p* < 0.05, \*\**p* < 0.01, \*\*\**p* < 0.001, \*\*\*\**p* < 0.0001. **e,** A schematic overview of lentiviral-based rat iPSC reprogramming and ChR2-eYFP insertion. After reprogramming, single-cell riPSCs were transduced, subcloned, and analyzed for expression of ChR2-eYFP. REF, rat embryonic fibroblast; riPSC, rat induced pluripotent stem cell; MEF, mouse embryonic fibroblast. **f,** Acute transverse slice from 2-4 wk old chimera brain generated with rat iPSCs engineered to express hChR2(H134R)-eYFP. eYFP expression was used to guide a pipette to eYFP-negative, mouse neurons for patch-clamp recordings. **g,** Evoked EPSPs in mouse neurons upon stimulation of rat neurons with blue light. Addition of glutamate receptor antagonists (CPP, NBQX) abolished light-evoked depolarizations indicating a postsynaptic response. Blue triangles mark blue light stimulation. Traces are an average of 20 trials. Scale is 2 mV, 50 ms. **h,** Rat neurons form functional synaptic connections with mouse neurons. The peak EPSP amplitude was blocked by glutamate receptor antagonists and occurred 5-15 ms following light stimulation. n = 10 cells, 7 animals. Significance was tested by a two-tailed, paired t-test. \*\**p* < 0.01.

Analyses of three cortical regions at three timepoints showed the same striking result; in all regions and timepoints, rat neurons developing in the mouse brain appear to adopt the developmental timing of the host, as shown by intermixing of mouse and rat cells in each layer (Fig. 2b, c), and co-expression of the layer IV and V marker CTIP2 (Bcl11b) (Supplementary Fig. 2a). To examine other brain regions, we used double labeling with BrdU at E12.5 and EdU at either E13.4 or 14.5 to examine three other brain regions (Hippocampus CA1, Piriform Cortex layer 2 and Caudoputamen). As in the cortex, rat cell birthdates matched those of the mouse in the hippocampus and piriform cortex, although we detected a very small but statistically significant reduction in the earliest born rat neurons in the subpallial region (Caudoputamen) (Fig. 2d, Supplementary Fig. 2a) This is consistent with the lower overall contribution of rat cells to subpallial regions in the whole brain imaging studies (Fig. 1f). Together these results illustrate that developmental timing of xenogeneic donor PSC-derived neurons can be reprogrammed, perhaps by non-cell-autonomous mechanisms specific to the host. This is reminiscent of results from other interspecies chimeras in which the host environment governs the body and organ size, as well as the development of a chimeric organ absent from the donor species (gall bladder)^3,5,7^.

## Rat neurons synapse with mouse neurons

Neurons relay information through synaptic neurotransmission. However, neurons from different species may differ in proteins or signaling molecules necessary to recognize and form connections with synaptic partners *in vivo*^21^. To determine whether rat neurons can synaptically communicate with mouse neurons after developing within chimeras, we generated a rat iPSC line expressing the hChR2(H134R)-eYFP variant of channelrhodopsin^22^ (riPSC::hSyn-ChR2-eYFP) (Fig. 3a, Supplementary Fig. 3a, b). This iPSC reporter line provides us a tool to visually identify rat neurons, selectively activate them using blue light in derived chimeras, and record synaptic responses in targeted neurons. Multiple chimeras with rat contributions in cortical and hippocampal regions were readily produced from hSyn-ChR2-eYFP iPSCs (Table 1). To detect interspecies synapses, we collected acute slices containing the hippocampus and cortex for electrophysiology recordings (Fig 3b). Recordings from mouse hippocampal and cortical pyramidal neurons (based on their lack of eYFP expression) revealed EPSPs in mouse neurons 5-15 ms after light-stimulation that were blocked by AMPA and NMDA glutamate receptor antagonists (Fig. 3c, d, Supplementary Fig. 3c). In contrast, recording directly from a rat neuron showed faster responses that were not blocked pharmacologically (Fig. 3d). Together, these findings indicate rat neurons can form functional synapses with mouse neurons in at least two brain regions of rat-mouse chimeras.

**Figure 3:**
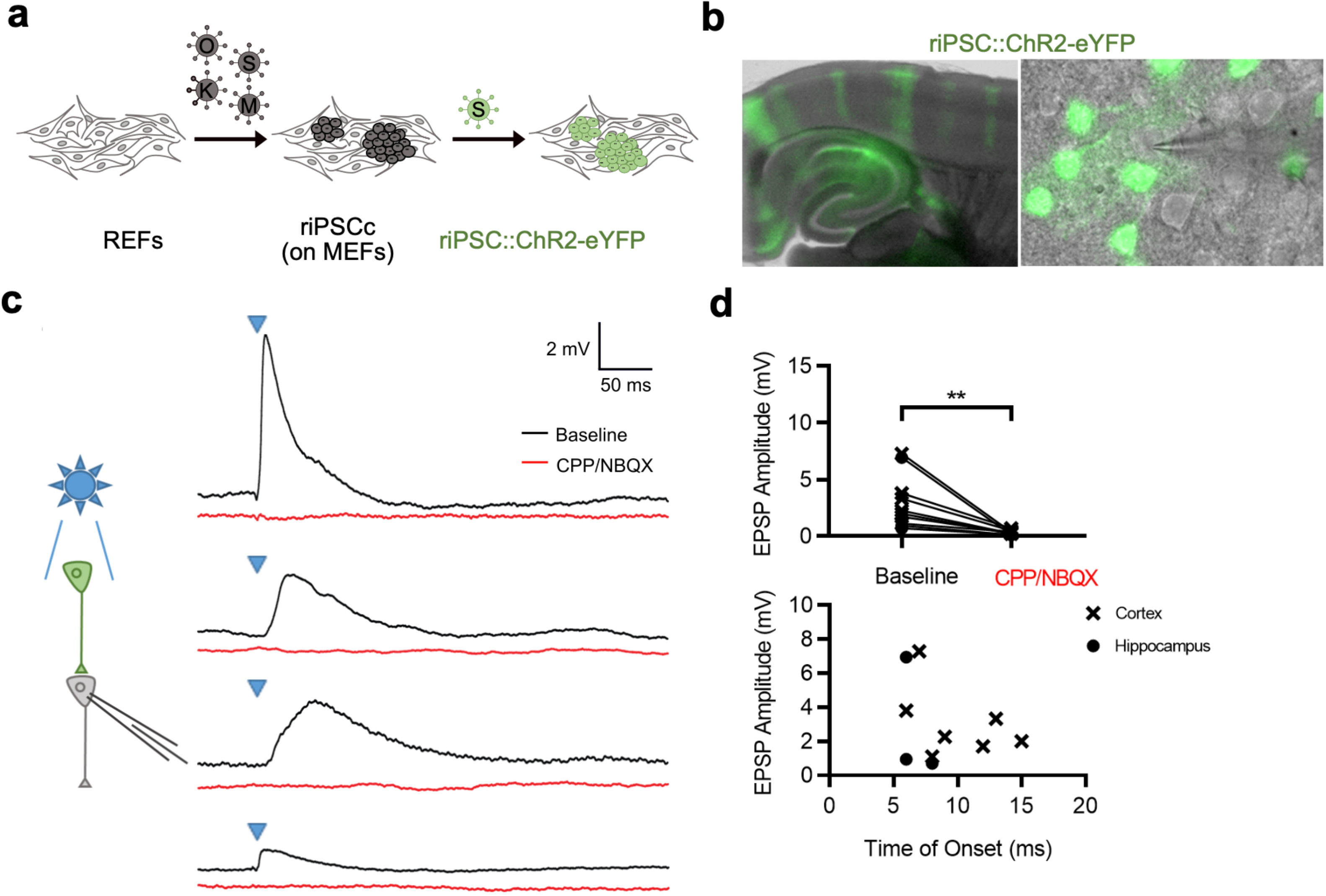
Rat neurons form functional synaptic connections with mouse neurons. **a,** A schematic overview of lentiviral-based rat iPSC reprogramming and ChR2-eYFP insertion. After reprogramming, single-cell riPSCs were transduced, subcloned, and analyzed for expression of ChR2-eYFP. REF, rat embryonic fibroblast; riPSC, rat induced pluripotent stem cell; MEF, mouse embryonic fibroblast. **b,** Acute transverse slice from 2-4 wk old chimera brain generated with rat iPSCs engineered to express hChR2(H134R)-eYFP. eYFP expression was used to guide a pipette to eYFP-negative, mouse neurons for patch-clamp recordings. **c,** Evoked EPSPs in mouse neurons upon stimulation of rat neurons with blue light. Addition of glutamate receptor antagonists (CPP, NBQX) abolished light-evoked depolarizations indicating a postsynaptic response. Blue triangles mark blue light stimulation. Traces are an average of 20 trials. Scale is 2 mV, 50 ms. **d,** The peak EPSP amplitude was blocked by glutamate receptor antagonists and occurred 5-15 ms following light stimulation. n = 10 cells, 7 animals. Significance was tested by a two-tailed, paired t-test. \*\**p* < 0.01.

## Intraspecies circuit rescue in genetic models of sensory dysfunction

Neural blastocyst complementation aims to engineer systems in which specific neuronal circuits can be rescued through selective cell replacement. Similarly, human neural cell replacement therapies are envisioned for cases where neurons are missing due to injury or cell death, and also when they remain present but have lost function or synaptic connectivity. Here, we test whether rat neurons can functionally complement two genetic mouse models of neural cell loss and neural cell synaptic impairment using the well characterized and genetically tractable circuits of the olfactory system.

Olfactory circuits begin in the olfactory epithelium (OE) of the nose, where olfactory sensory neurons (OSNs) detect odors. OSNs are produced by basal stem cells in the OE throughout the life of an organism and are one of the few sites of adult neurogenesis (Fig. 4a). Mature OSNs target their axons to one of two spatially invariant glomeruli in the olfactory bulb (OB) depending on their stochastic choice of one of ∼1200 genetically encoded olfactory receptors (ORs) (Fig. 4a)^23,24^. The general spatial position of glomeruli is conserved and similar between mouse and rat^25–27^, but the rat genome encodes at least ∼100 more ORs than the mouse, creating a larger OB with more glomeruli than in the mouse^28^. Finally, olfactory information travels to cortical processing centers via the olfactory mitral and tufted neurons (MT). By using the olfactory system, we seek to study how well blastocyst complementation can enable neurogenesis during throughout life, build anatomically recognizable circuit structures, permit appropriate axon guidance and synapse formation and determine whether exogenous neurons rescue local and distal circuit activity and/or olfactory behavior.

**Figure 4:**
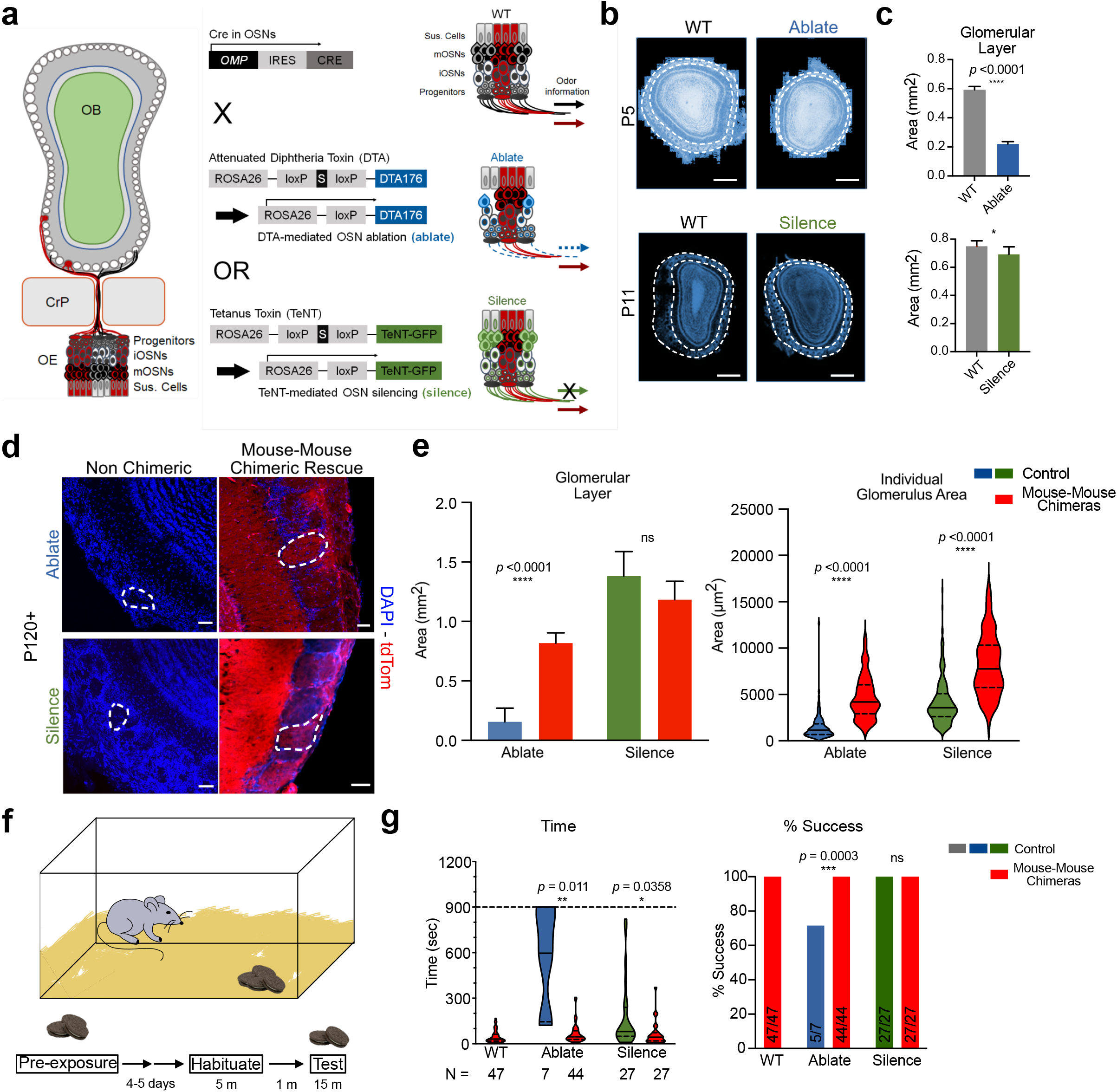
Genetic mouse models to test intraspecies olfactory complementation. **a,** Schematic of genetic strategies: crossing OMP-Cre mice with floxed DTA and TeNT animals results in recombination to either ablate (DTA) or silence (TeNT) mouse OSNs. **b,** Representative images of the olfactory bulb from wild-type (WT) mice and DTA-mediated OSN ablation (ablate) at P5 (top) (nuclei shown in blue, DAPI). Also shown are the olfactory bulbs in wild-type (WT) and TeNT-mediated OSN silencing (silence) at P11 (bottom) (nuclei shown in blue, DAPI). The region between the two dotted lines represents the glomerular layer. Scale bar 500 μm. **c,** Glomerular layer size is strongly reduced in the Ablate model and weakly in the silence model. Data shown are the means ± 95% CI; Silence; wild-type (WT): n = 9, Silence n = 6, Ablate; wild-type (WT): n = 15, Ablate n = 11. Significance was tested by *a* two tailed, unpaired t-test, \**p* < 0.05, **** *p* < 0.0001. **d,** Red mouse PSCs expressing tdTomato can rescue OB deficits in both models. Representative images from ablate (top) from non-chimeric (left) and mouse-mouse chimeric brain (right) rescue and from silence (bottom) from non-chimeric (left) and mouse-mouse chimeric brain (right) rescue are shown. Dotted lines indicate an individual glomerulus. Shown are the DAPI (blue) and tdTomato (red) channels. Scale bar 50 μm. **e,** Rescue of the area of glomerular layer (bar graph, left) and individual glomerulus size (violin plot, right) in ablated and silenced controls and chimeric rescues are shown. For glomerular layer size, data shown are the means ± 95% CI; Silence; control: n = 6, mouse chimera: n = 6; Ablate; control: n = 4, mouse chimera: n = 10. *P*-values shown are two-tailed unpaired t-test. \*\*\*\**p* < 0.0001. For individual glomerular size, violin plots with the median, upper and lower quartiles are plotted; Silence; control: n = 260 glomeruli, mouse chimera: n = 134 glomeruli; Ablate; control: n = 141 glomeruli, mouse chimera: n = 260 glomeruli. Significance was tested by two-tailed unpaired t-test, \*\*\*\**p* < 0.0001. **f,** Schematic of the buried cookie test. **g,** WT Mouse PSCs rescue behavior in both models as measured by the time to find the cookie and the success rates. For time, violin plots with the median and the upper and lower quartile are shown. WT (mouse mouse chimera): n = 47. Ablate (control): n = 7, Ablate (mouse rescue chimera) n = 44, Silence: n = 27, Silence (mouse rescue chimera) n = 27. Significance was tested by repeated measures One-Way ANOVA with Dunnes multiple comparisons test. For percent success, the test was deemed successful if the cookie was found under 900 seconds. For ablate control, 5 from 7 were successful with 44 from 44 rescued by mouse. For silence control, 40 from103 were successful with 27 from 27 rescued. Significance was tested by Fisher’s exact test, \*\**p<*0.01, \**p* < 0.05.

To this end, we have established two distinct mouse models in which OSNs are genetically disabled, then assess their compatibility with intra- and inter-species neural blastocyst complementation with mouse and rat PSCs, respectively. In mice expressing Cre recombinase under the control of the endogenous OMP locus, Cre is expressed highly specifically in mature OSNs, and has been widely used to mark and manipulate these cells ^27–29^. One model selectively ablates OSNs using Cre-activated expression of diptheria toxin subunit A (DTA, “Ablate” model) and a second model synaptically silences OSNs by selective expression of the tetanus neurotoxin light chain (TeNT, “Silence” model) ^27^. DTA expression in Ablate mice results in a loss of mature OSNs (Supplementary Fig. 4a, b) and a decreased glomerular layer area at P5 lacking in OSN axonal inputs to glomeruli in the OB (Supplementary Fig. 4c, d). In adult animals, however, a small number of misshapen glomeruli emerge (Supplementary Fig. 4c, d). In the second, “Silence”, model expression of TeNT in OSNs prevents neurons from releasing synaptic vesicles by cleaving VAMP, but allows normal OB formation as reported previously and shown here (Fig. 4a-c)^27,30,31^.

Olfactory behavior has not been tested using intraspecies (mouse-mouse) neural blastocyst complementation. In both the Silence and Ablate models, red (tdTomato-labeled) WT mouse PSC-derived neurons occupied the OE and projected axons to the OB. The donor mouse cells also rescued the anatomic aberrations in the OB of the Ablate mice (Fig. 4d, e). To test whether intraspecies olfactory complementation could rescue a primal olfactory behavior, food seeking, we employed the buried cookie test^32^. In this widely used assay, animals are habituated to a food reward, rested for 4-5 days, and video monitoring is used to score the time to find a hidden cookie (Fig. 4f). The poor success rates in olfactory guided food search in both the Ablate and Silence mouse models were rescued significantly by the donor WT mouse cells (Fig. 3g, h and Supplementary Fig. 4e). These results show that selective rescue of sensory cells and olfactory behavior is highly effective using intraspecies neural blastocyst complementation.

## Mouse-rat olfactory systems

Next, we aimed to use these two complementation systems to test the ability of rat neurons to support and participate in the formation and function of olfactory circuits in a mouse brain. First, we examined integration of rat neurons in the WT mouse olfactory system. We found that rat cells can differentiate into mature OSNs, based on their expression of OMP, appropriate laminar position and columnar structure in the OE and their ability to extend axons that navigate from the nose to innervate the olfactory bulb where they form distinct red rat glomeruli that can persist for ∼2 years (Fig. 5a, b, Supplementary Fig. 5a). Surprisingly, none of the mouse or rat glomerular structures in the chimeric brains showed innervation by rat mitral/tufted (MT) neuronal dendrites, the synaptic targets for OSNs, and we could not identify any rat neurons that resembled MT neurons based on morphology or laminar position. Whether the lack of MT neurons is due to unknown interspecies developmental barriers, their earlier than usual birthdate (e9.5), or simply their low abundance remains to be tested. However, rat cells contributed to other more abundant neuronal subtypes in the OB (inhibitory GCs and periglomerular neurons) as defined by their position and characteristic morphologies. Moreover, we could structurally demonstrate the formation of rat-mouse synapses in OB glomeruli using electron microscopy (Fig 5b, Supplementary Fig. 5b). Together these results establish the olfactory system as suitable for testing functional complementation in rat-mouse chimeras.

**Figure 5:**
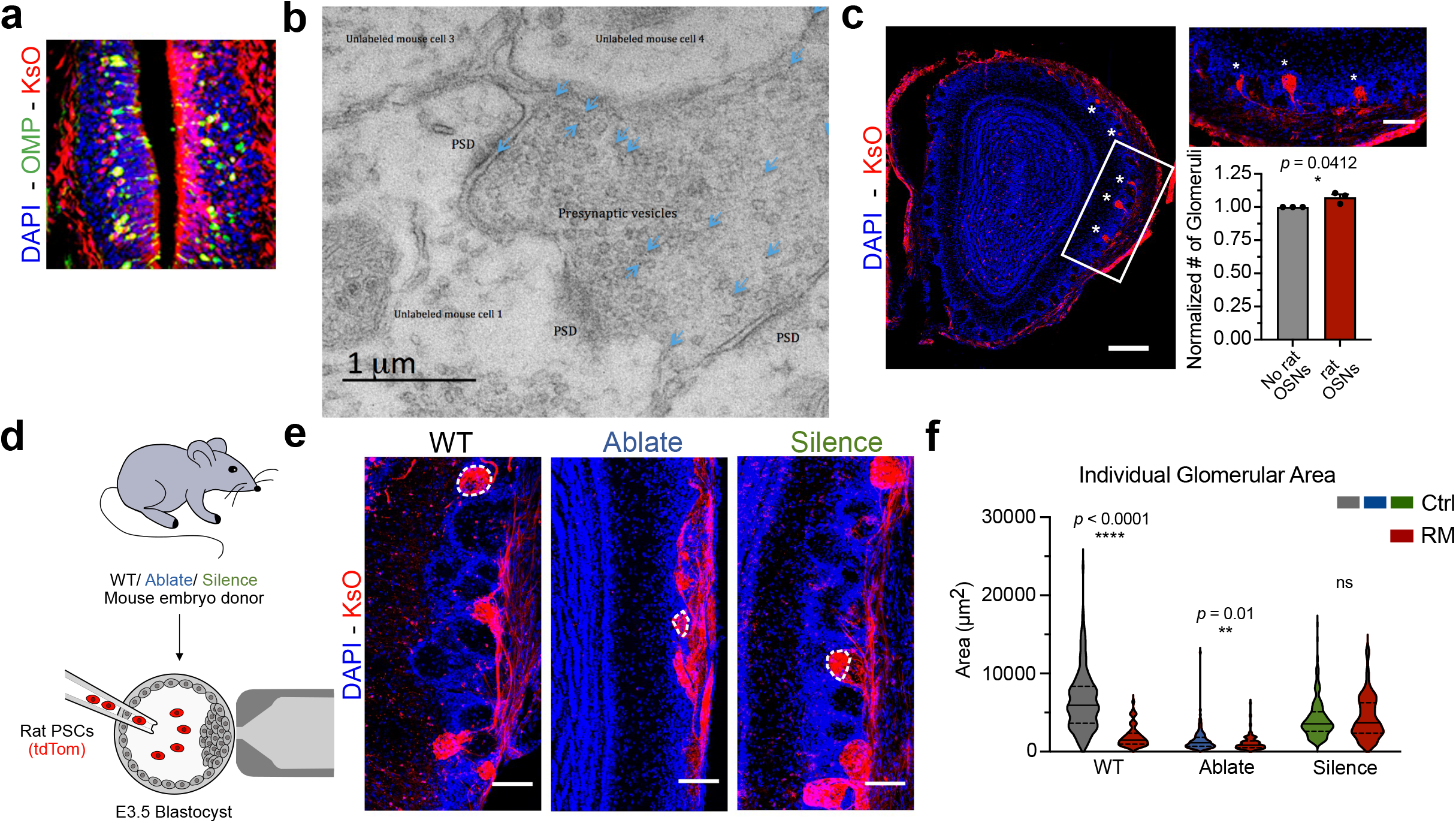
Rat-mouse chimeras complementation of olfactory system anatomy. **a,** A representative image of OMP-positive mature rat OSNs in a rat-mouse chimera. Shown are nuclei (DAPI, blue) OMP (green) and rat KsO (red). Scale bar, 100 μm. **b,** A representative image of transmission electron microscopy performed on rat-mouse chimeric synapses. The sections were labelled with immunogold staining against KsO as indicated by dark densities marked by blue arrows. Synapses formed between the rat and mouse cells with presynaptic vesicles and post-synaptic densities (PSD) are labeled. **c,** A representative serial OB section demonstrates the formation of glomeruli with majority rat cells or no rat cells in a P10 rat-mouse chimera. Shown are rat KsO (red) and nuclei (DAPI, blue). Asterisks mark rat glomeruli. White box indicates inset. Scale bars, 250 μm and 100 μm (inset). Quantification of the total number of glomeruli in animals with unilateral rat OSN contribution (right). The number was normalized to the hemisphere without rat glomeruli. Data shown are mean ± 95% CI, n = 3 animals, 32 slices/animal. *Significance was tested by* unpaired, two-tailed t-test. \**P* = 0.0412. **d,** Schematic of wild-type rat PSCs expressing KsO injected into mouse embryos from WT, Ablate and Silence models. **e,** Rat glomeruli (red) in chimeras generated from each transgenic cross, nuclei in blue (DAPI) outline structures. Dotted lines outline glomeruli. Scale bar, 100 μm. **f,** Rat vs. mouse glomerulus size across models. Violin plots with the median and upper quartile and lower quartile are shown; n = 3-4 animals/genotype, 2-3 slices/animal, 191 glomeruli (WT-mouse), 68 (WT-rat), 141 (ablate-mouse), 145 (ablate-rat), 260 (silence-mouse), 232 (silence-rat). Significance was tested by *2*-way ANOVA followed by Tukey’s multiple comparisons test. \*\*\*\**P* < 0.0001, \*\**P* < 0.01

Which genome might govern the formation of the olfactory sensory map in interspecies chimeras? In the intraspecies mouse-mouse glomeruli, donor red fibers intermix with the unlabeled host mouse axons (Fig. 4d). However, in mice genetically engineered to express a rat OR in place of a mouse receptor, mouse OSNs with the rat OR form unique glomeruli^25,26^. Here, we observe distinct rat and mouse glomeruli without apparent rat innervation, as in the transgenic mice (Figure 5c, Supplementary Fig. 5a). Because the rat OR repertoire is larger than that of the mouse, one might expect interspecies chimera to have more overall glomeruli. We counted the numbers of glomeruli from serial OB sections of young animals with rat OSN contribution to only one OB hemisphere (Fig. 5c). OBs with rat glomeruli had a small but significant increase in the number of glomeruli compared to OBs lacking rat OSN innervation, further suggesting that rat OSNs form additional independent glomeruli in the mouse OB (Fig. 5c). Overall, these results show that rat neurons can alter their birthdates and precise patterns (and lengths) of axonal projections based on cues in the mouse. Further, they may be capable of building an expanded olfactory detection repertoire using the increase in glomeruli and perhaps functionally distinct rat ORs.

## Anatomic and functional complementation in primary rat-mouse neural circuits

Having demonstrated that rat cells can become OSNs that exhibit apparently normal wiring to the mouse OB, we next wished to dissect how well they can rescue olfactory circuits in the models of neuron loss or synaptic dysfunction using the Ablate and Silence models. We tested this by injecting rat DAC2 ESCs into Ablate and Silence mouse blastocysts to produce WT rat-mutant mouse chimeras (Table 1, Fig. 5d). In Ablate animals with rat OSN contribution, rat glomeruli were significantly smaller and more disorganized than those found in WT mice (Fig. 5e, f). In contrast, Silence model chimeras exhibited robust innervation of glomeruli and a largely normal glomerular area (Fig. 5e, f). One explanation for these observations may be that rat OSNs require the structure and cues, but not neural activity, from mouse OSNs to navigate to the OB and form robust glomeruli.

To assess functional rescue, we first assessed local activity in both models. In the OB, inhibitory periglomerular neurons surrounding glomeruli receive direct input from OSNs and express tyrosine hydroxylase (TH) in an activity-dependent manner^33^. We could detect weak rat driven TH expression in periglomerular neurons in the WT mouse, but not Ablate, chimeras (Fig. 6a, b,). In contrast, rat glomeruli in Silence chimeras had significantly higher TH expression than neighboring mouse glomeruli in the same OB, indicating that they may be “winners” in the known activity dependent competition between glomeruli in the OB (Fig. 6a, b). These results show that rat OSNs selectively elicit activity in local mouse interneurons, demonstrating appropriate synaptic connectivity and activation, similar to that of mouse OSNs.

**Figure 6:**
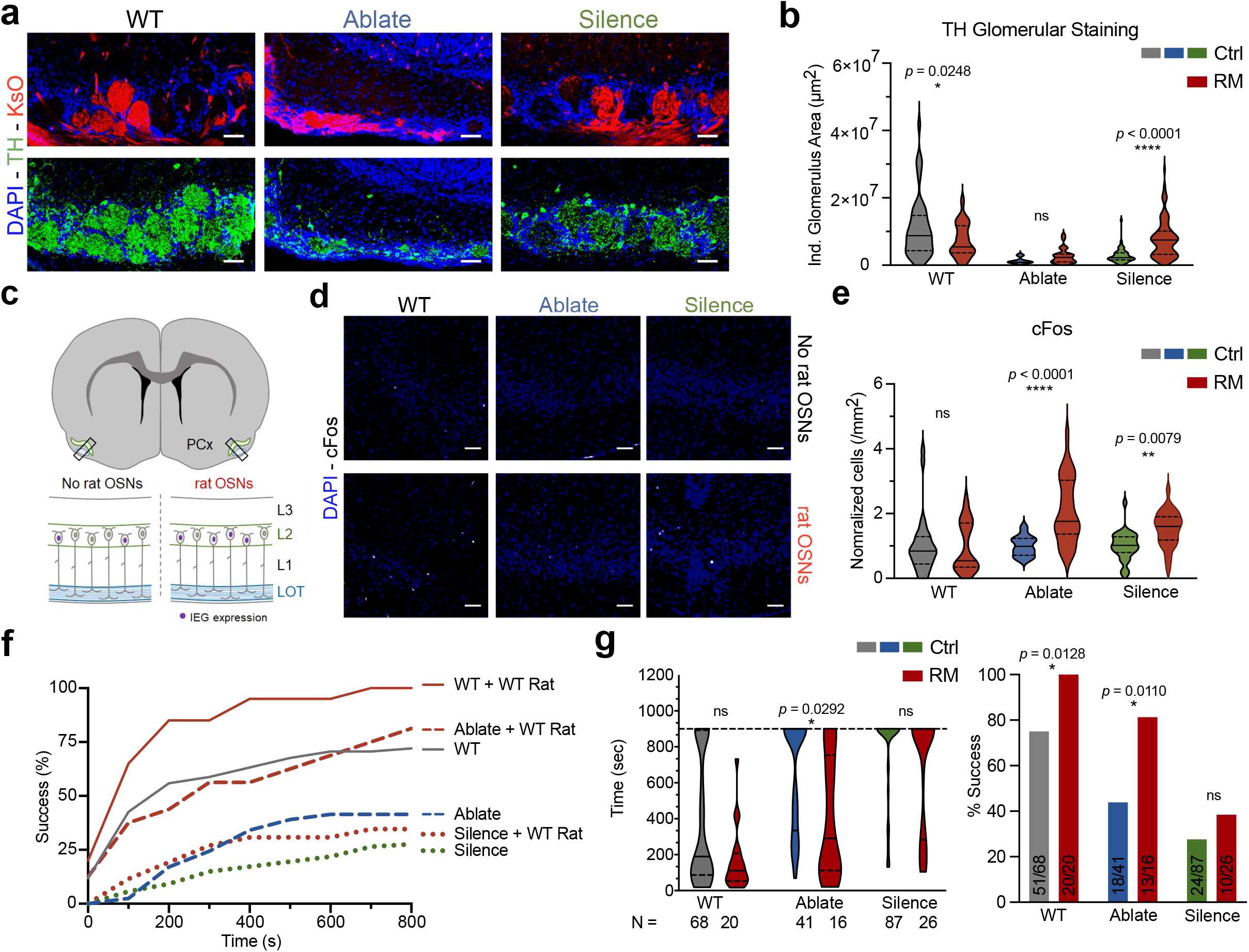
Intraspecies chimeras rescue olfactory behavior. **a,** Rat and mouse glomeruli stained for tyrosine hydroxylase (TH, green), nuclei (DAPI, blue), rat KsO (red). Scale bar, 50 μm. **b,** Rat glomeruli rescue TH only in the silence model. Violin plots with median and upper and lower quartiles are shown; n = 1 animal/genotype, 3 slices/animal, 44 glomeruli (WT-mouse), 23 (WT-rat), 5 (ablate-mouse), 25 (ablate-rat), 60 (silence-mouse), 95 (silence-rat). Significance was tested byTwo-way ANOVA followed by Sidak’s multiple comparisons test. \**p* < 0.0248, \*\*\*\**p <* 0.0001. **c,** A schematic illustrating the location of the neurons analyzed for the expression of immediate early genes (IEGs) in the parietal cortex of animals with unilateral rat OSN contributions. **d,** Representative images of cFos in PCx of animals with unilateral OSN contribution. Shown are the nuclei (DAPI, blue) and cFos (white). Scale bar, 50 μm. **e,** Rat OSN contribution increases cFos expression in the ipsilateral PCx of OSN-compromised animals. Cell densities were normalized to the mean density in the no contribution hemisphere for each animal. Violin plots with the median and upper and lower quartiles are shown; n = 2 animal/genotype, 15 slices/animal. *Significance was tested by* two-way ANOVA followed by Sidak’s multiple comparisons test, \*\**p* < 0.01, \*\*\*\**p*< 0.0001. **f,** WT rat PSCs rescue behaviour in both models of affected olfaction in mice. Shown are the percentage of the mice that find the cookie, plotted as success percentage against time (s). **g,** Selective rescue of olfaction in the ablate but not silence model. For time, violin plots with the median and the upper and lower quartile are shown; Wild-type: n =68, Wild-type (rat-mouse chimeras): n = 20, Ablate (control): n = 41, Ablate (rat-mouse chimeras): n = 16, Silence (control): n = 87, Silence (rat-mouse chimeras): n = 26. Rat cells influenced time to find the cookie in Ablate (p=0.0292) but not the Silence model. Significance was tested by repeated measures One-Way ANOVA with Dunnes multiple comparisons test. Success rates show the same trend with 900s as a cutoff. For WT, 51/68 trials succeeded. For WT Rat-Mouse chimeras, 20/20 trials succeeded (p=0.0128). For ablate models, 18/41 succeeded but with Rat-Mouse chimerism, 13/16 succeeded (p=0.0110). For silence models 24/87 succeeded while 10/26 succeeded with contribution from rat cells (non-significant). Significance was tested by Fisher’s exact test, *p < 0.05.

## Cortical processing and olfactory behavior in distinct rat complementation models

Rat OSNs can drive activity-dependent gene expression in the OB. However, the next stage of olfactory processing requires that OSNs form synapses with MT neurons, then correctly signal to the target neurons in the cortex. Rat OSNs might have innate differences in their synaptic strength, timing of vesicle release, or other characteristics that disrupt interspecies information transfer. To test this, we took advantage of the fact that olfactory information is transmitted to the cortex only ipsilaterally and not contralaterally^34–41^. In both our Ablate and Silence models, and in WT mice, we identified chimeras with unilateral rat contribution to OSNs in the OE and OB. We reasoned that we could determine whether rat neurons can communicate to the mouse olfactory cortex by directly comparing the silent “mouse-only” cortical hemisphere to the potentially rescued “rat-mouse” hemisphere, using serial sections take from the same animal’s brain (Fig. 6c). This design is important because the chimeras have stochastic contribution to OSNs, such that different chimeras are likely to have different OR repertoires. This could render them insensitive to certain individual odors. Measuring baseline activity in two hemispheres of the same brain across the same A-P axis in a complex odor environment should control for any unknown differences in odor stimuli between animals.

Here we analyze the piriform cortex (PCx), which is responsible for odor perception and discrimination^34–41^. Loss or silencing of OSNs, achieved by multiple methods, reduces activity and IEG expression in the piriform cortex (PCx), while sensory activity reliably increases c-Fos expression in the (PCx)^34–41^. Therefore, markers of neuronal activation in the PCx are well established as an indirect readout of OSN activity^38^. First, we examined c-Fos expression in the PCx of chimeras with unilateral OSN-contribution in WT, Ablate and Silence mouse models. Rat OSN contribution significantly increased cFos expression in the ipsilateral PCx of both Ablate and Silence chimeras (Fig. 6d, e, Supplementary Fig. 6a, b). This increase was not reflected in WT chimeras. We also tested expression of Egr1, which is less sensitive to direct odor stimulation and also has a higher background expression level ^39–41^. The natural odor environment appeared to be insufficient to evoke increased Egr1 expression, except for a very small increase in Silence chimeras (Supplementary Fig. 6c, d). Thus, these data demonstrate that sensory information flow derived from peripheral rat sensory neurons can be communicated to the PCx at the sensitivity level of c-Fos induction. Unexpectedly, the rescue of the Ablate model was greater than in the Silence model, even though the IEG (TH) activity in the local OB circuit, showed the reverse pattern. This was also puzzling because the Ablate OBs were misshapen and smaller than WT, even when rat OSNs were present. One explanation for this could be that rat OSN signals are stronger in the PCx due to the smaller OBs that might lead to less local inhibition, but many other explanations are possible.

Finally, we wished to test whether rat OSNs could enable an anosmic mouse to smell. We again employed the buried cookie test (see Methods). Unexpectedly, we show that rat cells can rescue this behavior in the Ablate, but not Silence models with respect to percent success, and time to find cookie, which are also plotted together as cumulative frequency distributions (Fig. 6f, g). This is consistent with the increase in cFos staining in the cortex of Ablate rat chimeras. However, neither this effect nor the rat rescue of the Silence mouse OSNs was as dramatic or complete as the mouse-mouse rescues shown in Fig. 4.

WT mouse olfaction appeared slightly improved in rat-mouse chimeras. However, this was not significant using a strict test for multiple comparisons (Dunn’s) but was significant using a test based on false discovery rates (Benjamini and Hochberg) (Supplementary Data Table 1). These results highlight the importance of testing different models of olfactory impairment across various aspects of neural circuit formation.

## Discussion

In conclusion, despite more than 10 million years of evolutionary divergence, when present from the earliest stages of mouse embryonic development, rat PSCs can produce neural cells that readily adapt to their host environment. Rat cells robustly specify into diverse morphological subtypes, build layered cortical structures synchronously with the host mouse cells, establish dynamic synaptic structures that persist for nearly two years and engage in multilevel synaptic communication from the periphery to the mouse cortex. Interestingly, despite this remarkable interspecies plasticity, we find that certain rat-specific programs overrule mouse developmental control. For example, rat neurons form OBs with an increased number of rat-specific glomeruli. Rat neurons also can fail to read important cues; for example, they cannot rescue OB formation in Ablate models.

Moreover, we show that while rat cells can partly rescue the primal mouse olfactory behavior of food seeking through selective complementation of its OSNs, they do not restore the sense of smell as well as intraspecies mouse complementation. This disparity might be explained somewhat trivially by the generally lower contribution of rat cells than we observe in intraspecies experiments. Alternatively, or in addition, there may be species-specific blockades to behavioral rescue. These could include differences in the molecules that drive the finer details of synaptic connectivity, set neuronal firing rates, or regulate intrinsic properties, such as excitability, that may have evolved differently across species with different sized brains. In support of this idea, mouse and rat proteins differ by about 4% at the amino acid level (93.6% conservation)^42^. This percent divergence can have clear consequences for circuit formation. For example, the OR I7 is 95% conserved between rat and mouse and responds to the same odor ligands^43^. Yet transgenic expression of rat OR I7 in mouse OSNs leads to formation of an independent rat glomerulus that does not mix with the mouse^44^. Related to this, a unique strength of the blastocyst interspecies chimera over transplant models is that it allows rat neurons to transit through all stages of embryonic and brain development concomitantly with the mouse. This exposes the rat cells to multiple levels of cell-cell competition, in which they might fail to wire correctly. Although this may be rescued if the mouse homologous cell type is absent or disabled, as we report here. Interspecies brain chimeras, therefore, offer a unique tool to explore the molecular basis for these barriers.

Accordingly, in the behavior rescue experiments, we report several interesting, and somewhat unexpected results. First, it was surprising to us that there was weak but detectable improvement in performance of WT animals complemented with rat cells. Second, it was surprising to us that the Ablate models performed better than the Silence models, despite their aberrant OB structures and local circuits. One model to explain this could be that having more active glomeruli improves olfaction. In the WT mice, perhaps the supernumerary rat glomeruli provide new sensory can modestly improve olfaction in a WT mouse and supply sufficient signal to provide rudimentary signaling in otherwise silent cortical circuits. Yet, in the Silence model, the presence of weakly active OSNs could provide some sort of competition that either dampens or de-correlates signaling from the rat, and thus scrambles its olfactory responses. Absent the “noise”, perhaps the few rat glomeruli in Ablate models have a more direct path to activate the cortex sufficiently well to provoke food seeking. Dissecting these intriguing questions would benefit from means to drive rat neurons more exclusively towards the desired cell fates.

Despite current technical limitations due to stochastic contributions in chimeras, through our use of clean genetic systems and large numbers of chimeras, we have provided proof-of-principle that interspecies neural blastocyst complementation can be a powerful tool. Extending this tool to different genetic models and more distant species may provide novel discoveries about brain evolution and help build better models of human neurologic disease and brain repair therapies.

## Methods

### Mouse strains

CD1 animals were used for surrogate mothers and to generate MEF feeders. Omp-IRES-Cre^29^ mice were generated by Kristin Baldwin in the Axel laboratory and crossed with C57BL/6J mice to generate heterozygotes. The TeNT-GFP mouse strain (TeNT) was previously provided by Martin Goulding^45^ and ROSA-DTA (DTA) mice were ordered from The Jackson Laboratory (Stock # 009669)^46^. Omp-IRES-Cre x C57BL/6J F_1_ mice were bred with either TeNT or DTA mice to generate transgenic blastocysts and internal wild-type littermate controls. DBA/2J mice were crossed with C57BL/6J mice to generate C57BL/6J x DBA F_1_ males for WT blastocyst generation and resulting chimeras were used for electrophysiology and birth dating experiments^47^. Both females and males were used for all experiments. All animal experiments were conducted in accordance with the protocols approved by the IACUC of The Scripps Research Institute and NIH guidelines for animal use.

### Genomic DNA extraction and PCR

Genomic DNA was extracted from mouse tail tip samples using the REDExtract-N-Amp Tissue PCR kit (Sigma-Aldrich). PCR was performed using the REDExtract-N-Amp PCR Readymix (Sigma-Aldrich). The following primers were used for genotyping: *Omp* WT Sense: 5’– TGTATTTCCTCATCACCTTTGGCG–3’; *Omp* WT Antisense: 5’– GGTCAGTCTCTTATCTCTCAGTCCCG–3’; Cre Sense: 5’–CGCATAACCAGTGAAACAGCA–3’; Cre Antisense: 5’–CGCATAACCAGTGAAACAGCA–3’.

For species genotyping and validating hSyn-ChR2-eYFP subclones, genomic DNA was extracted from rat PSC cell pellets using the DNeasy Blood and Tissue kit (Qiagen) and PCR was performed using the Platinum PCR SuperMix High-Fidelity kit (Invitrogen). Rat PSC lines were validated by genotyping for the rat specific *Omp* allele using the following primers: *Omp* Mouse Sense: 5’–CCTGACAGGGGCTATGACAGAGTG–3’; *Omp* Rat Sense: 5’–GGCAGTATGCGGTTGGATCAATCAG–3’; *Omp* Common Antisense: 5’– CCTGGTCCAGAACCAGCGGC–3’. To validate hSyn-ChR2-eYFP insertion within subclones, the following primer sets were used for ChR2: ChR2 Sense 1: 5’–GGATTGAATCTCGCGGCACG– 3’; ChR2 Antisense 1: 5’–GTTGCCATGGCGCTGGTAGC–3’; ChR2 Sense 2: 5’– GCGTCCTGAGCGTCTATGGC–3’; ChR2 Antisense 2: 5’–GCTTGCCGGTGGTGCAGATG–3’; ChR2 Sense 3: 5’–GTTCATCTGCACCACCGGCA–3’; ChR2 Antisense 3: 5’– GCACGCTGCCGTCCTCGATG–3’.

### Lentiviral constructs and production

The Syn-hChR2(H134R)-eYFP lentiviral plasmid was ordered through Addgene (#20945)^22^. The rtTAM2.2 cassette (rtTA) was generated previously^48^ and cloned into a lentiviral transfer plasmid by the K. Baldwin laboratory^47^. Doxycycline (dox)-inducible lentiviral reprogramming factors encoding mouse cDNAs for Oct4, Sox2, c-Myc, and Klf4 were generated previously^47^. Virus was produced in HEK293T cells (ATCC CRL-3216) by calcium phosphate co-transfection of lentiviral shuttle and packaging vectors (pRRE (Addgene #12251), pRev (Addgene #12253), pMD2.G (Addgene #12259))^49,50^. Live lentivirus was harvested 48 hr. post-transfection for transduction.

### Rat iPSC reprogramming

Rat embryonic fibroblasts (REFs) from WT Sprague-Dawley rats were obtained from ATCC (CRL-1213) and reprogrammed into iPSCs using dox-inducible lentiviral vectors according to previously established protocols^3,47^. New rat iPSC lines were validated by immunocytochemistry for pluripotency markers (Sox2, Oct4, SSEA1, Nanog).

### Lentiviral-integrated iPSC subclones

To generate riPSC::hSyn-ChR2-eYFP lines, iPSCs were separated from mitotically inactive MEF feeders by culturing on Matrigel-coated (Corning) tissue culture flasks for 3 passages. hSyn-hChR2(H134R)-eYFP lentivirus was added to single cells after the third passage and left overnight. The following day, lentivirus was removed, iPSCs were washed twice with phosphate-buffered saline (PBS), passaged, and then seeded at 5×10^5^ cells on fresh feeders in 10-cm^2^ tissue culture dishes. Three days after seeding, subclones were manually picked using fluorescence as a guide, expanded, and validated for successful ChR2 integration by PCR.

### Cell culture conditions

HEK293T cells, MEFs, and REFs were cultured in standard media (DMEM with GlutaMAX (Gibco), 10% heat-inactivated fetal bovine serum (Gemini Bio), 1X penicillin-streptomycin (Gibco)). MEF feeders were inactivated at passage 4 with chemical treatment of 1 μg/ml mitomycin-c (STEMCELL Technologies) overnight. Rat PSCs were cultured on mitotically inactivated MEFs in serum-free 2i media ((N2B27 basal medium [1:1 DMEM/F12 (Gibco), N2 supplement (Gibco): Neurobasal A (Gibco), B27 supplement minus vitamin A (Gibco)), 100 μM β-mercaptoethanol (Gibco], supplemented with 10 ng/mL mouse LIF (STEMCELL Technologies), 3 μM CHIR99021 (STEMCELL Technologies), 1 μM PD0325901 (STEMCELL Technologies)) and passaged as single cells onto fresh feeders using TrypLE Express (Gibco). All cells were kept at 37°C in a humidified environment at 5% CO_2_.

### Generation of chimeras

DAC2 and DAC8 cells lines were provided by the Ying laboratory^3,12^. SDFE and SDFF rat iPSC lines were previously reported^3^. SDMJ rat iPSC lines were generated by the Belmonte lab and SD riPSC 9.3 by the Baldwin lab. The mouse PSC line (31-2/B2) was previously generated by the Baldwin lab^51^. Chimeras were produced by injection of PSC cells into E3.5 blastocysts, collected from superovulated C57BL/6J females mated to C57BL/6J x DBA2 F_1_ stud males, as previously reported^3,44,48,^.

### Tissue preparation, immunohistochemistry (IHC) and immunocytochemistry (ICC)

Mice P21 were euthanized with isoflurane and transcardially perfused with 4% PFA (EMS). Brains with OBs attached and the OE were dissected from the skull and placed in 4% PFA solution overnight at 4°C. For mice P10 and younger, animals were briefly anesthetized, decapitated, and the entire head fixed in 4% PFA solution overnight at 4°C. After post-fixation, brains were washed in PBS. Cells in culture were washed and then fixed in 4% PFA for 10 min at room temperature.

For vibratome sections, brains were embedded in 4% low-melt agarose (Bio-Rad) and sectioned in the coronal plane at 80 μm using a Leica VT1000S. For cryostat sections, samples were submerged in 30% sucrose (Acros Organics) at 4°C and left overnight or until the tissue sank to the bottom of the tube. OE samples from mice P21 and older were incubated in 0.5 M EDTA solution, pH 8.0 (Invitrogen) for 1 hr. at room temperature to decalcify before dehydrating in 30% sucrose. Dehydrated samples were embedded in Tissue-Tek O.C.T. (Sakura), rapidly frozen in a bath of 70% ethanol and dry ice, and stored at -80 °C. Before sectioning, samples were moved to -20°C for 1 hr. Using a Leica CM3050 S, 30 μm cryosections were collected and allowed to air dry for 1 hr. before staining or storage at -80°C.

For IHC and ICC, vibratome or cell culture samples were blocked in 10% heat-inactivated horse serum (Lonza BioWhittaker) and 0.1% Triton X-100 (Sigma-Aldrich) for 1 hr. at room temperature. Primary antibodies were diluted 1:500 (IHC) or 1:250 (ICC) in blocking buffer and added overnight at 4°C. Sections were washed three times and then stained with secondary antibodies and DAPI (5 mg/ml), both diluted 1:1000 in 0.1% Triton X-100, for 1 hr. at room temperature. Cryosections followed the same protocol with a few additions. Slices were removed from -80°C and brought to room temperature for ∼30 min. In cases of antigen retrieval (AR), slides were submerged in sodium citrate buffer (Sigma Aldrich) warmed to ∼85-90°C on a hot plate for 20 min. All sections were washed twice to rehydrate samples and remove residual O.C.T. before blocking.

The following primary antibodies were used: KsO (MBL PM051M, Rabbit, Polyclonal, IHC), BrdU (BD Pharmingen 555627, Mouse, Monoclonal, IHC – HCl treatment), Oct4 (SCBT sc-5279, Mouse, Monoclonal, ICC), Sox2 (SCBT sc-365823, Mouse, Monoclonal, ICC), SSEA1 (SCBT sc-21702, Mouse, Monoclonal, ICC), Nanog (R&D Systems AF2729, Goat, Polyclonal, ICC), GFP (Invitrogen A10262, Chicken, Polyclonal, ICC – 1:500), OMP (Wako 544-10001, Goat, Polyclonal, IHC – AR), TH (Pel-Freez P40101-150, Rabbit, Polyclonal, IHC), cFos (CST 2250S, Rabbit, Monoclonal, IHC), and Egr1 (SCBT sc-189, Rabbit, Polyclonal, IHC). The following secondary antibodies were used: Alexa Fluor 488 (Abcam ab150169, Goat anti-Chicken IgY H&L), Alexa Fluor 488 (Invitrogen A-21202, Donkey anti-Mouse IgG H&L), BP-FITC (SCBT sc-516140, anti-Mouse IgGκ), Alexa Fluor 488 (Invitrogen A-21206, Donkey anti-Rabbit IgG H&L), Alexa Fluor 555 (Invitrogen A-31570, Donkey anti-Mouse IgG H&L), Alexa Fluor 555 (Invitrogen A-31570, Donkey anti-Mouse IgG H&L), Alexa Fluor 555 (Invitrogen A-31572, Donkey anti-Rabbit IgG H&L), and Alexa Fluor 647 (Abcam ab150131, Donkey anti-Goat IgG H&L).

### Whole-brain image acquisition

Animals were perfused, brains post-fixed overnight with 4% PFA, and samples cleared using a water-based high-refractive index clearing solution (Zhuhao Wu; manuscript in preparation). The cleared brains were imaged on a commercial light-sheet microscope (Zeiss Z.1 Lightsheet), equipped with two 10× illumination objectives (0.2 NA) and a 5× detection objective (0.16 NA), at a pixel x-y-z resolution 1.26 × 1.26 × 8.04 μm. Every optical section was acquired as a tiled mosaic, with 488 nm and 561 nm channels scanned sequentially.

### Whole-brain registration

Imaged brains from 6-8 wk., mixed sex chimeras were registered to a standardized mouse reference STP (RSTP) brain as previously described^53–55^. Initial 3D affine transformation was calculated using 6 resolution levels, followed by a 3D B-spline transformation with 3 resolution levels. Similarity was computed using Advanced Mattes mutual Information metric by *Elastix* registration toolbox^56,57^. In order to enhance the precision of the image registration, both the acquired brain images and the reference brain were pre-processed used custom scripts to enhance anatomical landmark features used in the computation of mutual information (Muñoz Castañeda & Osten; manuscript in preparation).

### ROI Brain Area Analysis

Anatomical-based segmentation of the 3D registered brains was done using the 2011 Allen Reference Brain Atlas (ARA) labels with modifications as previously described. The developmental origin based segmentation was achieved using the Allen Developing Mouse Brain Atlas labels^49^ registered to the RSTP brain as described above for whole-brain registration, with the transformation parameters obtained from the standardized RSTP brain applied to warp and align the atlas onto the chimera brains.

### Brain Signal Segmentation

Whole brain signal distribution of the KsO-labeled rat cells was automatically detected using custom scripts. First, the signal background was reduced using a gamma correction filter and the signal threshold was set using Otsu’s algorithm; second, pixels detected were located in the 3D brain coordinates and mapped onto the anatomical brain areas of the reference atlases; third, the signal was quantified as the number of pixels per anatomical regions either in the standard ARA atlas or in the Allen Developing Mouse Brain Atlas; finally, the density analysis was expressed as the pixel volume per brain area volume for all anatomical regions as described previously^40,41^. All quantifications were done separately for each hemisphere.

### Confocal imaging, image processing, and analysis

All images were acquired on a Nikon A1 confocal microscope as large-image z-stacks. Analysis was performed using Nikon NIS-Elements. To count glomeruli, circular spaces were identified in the glomerular layer (GL) devoid of DAPI staining. For glomerular counts in P10 chimeras, glomeruli were manually counted across 32 serial sections and the total normalized to the hemisphere with no rat OSNs for each animal. To calculate the area of the GL, maximal coronal sections just anterior to the accessory olfactory bulb were used. Using DAPI counterstaining as a guide, the interior and outer edges of the GL (omitting the olfactory nerve layer) were traced, and the difference between these areas was calculated. For glomerular area in 7-14 wk. adult mice, borders were drawn for each glomerulus following DAPI and sorted by KsO expression. For TH intensity, glomeruli were outlined and the integrated density calculated using Fiji^59^. Cell counts (OMP, Egr1, cFos) were done manually in respective neuronal layers. The OE was defined using DAPI as the region between the lumen and lamina propria, and layer 2 of PCx as the dense band of nuclei. For IEG staining in PCx, serial sections were collected from the OB through PCx on the cryostat in order to maintain brain orientation. OB sections were visualized to identify animals with unilateral rat OSN contribution and to which hemisphere. Cortical sections were subsequently stained for cFos and Egr1, imaged, and cells manually counted within layer 2 of PCx. Cell densities were normalized within each slice to the hemisphere without rat OSN contribution. Statistical analyses were run using Prism 8 (GraphPad).

### Dual-pulse birth dating, staining, and analysis

Pregnant surrogates were given intraperitoneal injections delivering 50 mg/kg of BrdU (Life Technologies) at E12.5 and 50 mg/kg of EdU (Life Technologies) at either E13.5, E14.5, or E17.5 to label neurons in developing WT chimera embryos. Reconstituted as 10 mg/ml (BrdU) and 5 mg/ml (EdU) in dPBS, this equated to 200 μl and 400 μl injections, respectively, for a 40 g mouse. Chimeric brains were harvested at 4 wks. for IHC as described above.

For BrdU-EdU double staining, the following modifications were made to the IHC protocol. Slices were pre-stained with the KsO antibody in blocking buffer overnight at 4°C. Slices were washed three times, fixed again with 4% PFA for 15 min at room temperature, washed, and incubated in preheated 1N HCl solution for 30 min at 37°C. To neutralize, 0.1 M sodium borate buffer pH 8.5 was added at room temperature for 10 min. Slices were washed twice, blocked for 1 hr., and stained overnight at 4°C with BrdU and KsO primary antibodies. The subsequent day, slices were washed, stained with secondary antibodies, and then mounted onto slides to dry for 15 min at room temperature. The Click-iT EdU Cell Proliferation Kit for Imaging (Invitrogen) was used to identify EdU-positive cells following the manufacturer’s instructions starting with step 4.1 except 0.1% Triton X-100 was substituted for 3% BSA, slides were incubated with the reaction cocktail for 1 hr., and were stained for DAPI in 0.1% Triton X-100 for 1 hr.

To calculate the location of EdU-positive cortical cells, an arc was drawn to mark the border between cortex and corpus callosum (CC). Neurons were manually identified, sorted by KsO double staining, and their shortest distance to the marked border measured. For BrdU-EdU double staining, EdU-positive and BrdU-positive/EdU-negative labelled cells were manually identified and sorted by KsO expression. Cell densities were normalized to an average density of DAPI-stained nuclei (mouse) or the density of KsO-positive (rat) cells present in corresponding regions. Average DAPI densities were generated by counting the area ∼250 DAPI-stained nuclei occupy across 8 samples. Analysis was performed using R (3.5.1) and Prism 8 (GraphPad).

### Acute slice electrophysiology, recording, and analysis

WT chimeras were generated using riPSC::hSyn-ChR2-eYFP subclone #9.3 and used from 2-6 wks. of age for experiments. To isolate slices containing the Hipp and cortex, animals were anesthetized briefly with isoflurane then decapitated. The skin and skull were removed to expose the brain, which was rapidly dissected out and placed in ice-cold sucrose cutting solution (100 mM Sucrose (Sigma-Aldrich), 60 mM NaCl (Sigma-Aldrich), 26 mM NaHCO_3_ (Sigma-Aldrich), 20 mM D-Glucose (Sigma-Aldrich), 1.25 mM NaH_2_PO_4_-H_2_O (Sigma-Aldrich), 2.5 mM KCl (Honeywell Research Chemicals), 5 mM MgCl_2_-6H_2_O (Honeywell Research Chemicals), 1 mM CaCl_2_ (Honeywell Research Chemicals)) bubbled with 95% O_2_/5% CO_2_. Blocking cuts were made to remove the CB and OBs, bisect the hemispheres, and orient Hipp in the transverse plane. The blocked tissue was transferred to the slicing chamber of a Leica VT1200S vibratome containing cold, oxygenated sucrose cutting solution. Isolated 300 μm thick slices were transferred to a recovery chamber containing ACSF (127 mM NaCl, 25 mM NaHCO_3_, 25 mM D-Glucose, 1.25 mM NaH_2_PO_4_-H_2_O, 2.5 mM KCl, 2 mM CaCl_2_, 1 mM MgCl_2_-6H_2_O) bubbled with 95% O_2_/5% CO_2_ to recover for 30 min at 32°C, and were then maintained at room temperature.

Slices were transferred to a recording chamber and perfused with oxygenated ACSF warmed to 31°C. Whole-cell current clamp recordings were obtained from eYFP-positive rat and eYFP-negative mouse neurons visualized using a Scientifica SliceScope fitted with a SciCamPro for infrared differential interference contrast (IR-DIC) and fluorescence optics. Patch pipettes (3-5 mΩ) were filled with potassium gluconate internal solution (130 mM K-gluconate (Sigma-Aldrich), 11.5 mM Na_2_-phosphocreatine (Sigma-Aldrich), 10 mM HEPES (Sigma-Aldrich), 3 mM MgCl_2_, 3 mM Na_2_-ATP (Sigma-Aldrich), 0.2 mM Na-GTP (Sigma-Aldrich), 0.2 mM EGTA (Sigma-Aldrich), pH = 7.2, osmolarity = 307 mOsm). Current was applied to maintain cells at -70 mV and series resistance was monitored throughout recordings. Experiments were discarded if the holding current was greater than -300 pA, if the series resistance was greater than 25 MΩ, or if the series resistance changed by more than 20%. All recordings were acquired using a Multiclamp 700B amplifier and ScanImage software^60^. Signals were sampled at 10 kHz and filtered at 6 kHz. Analysis was performed using IGOR Pro (WaveMetrics).

Light-evoked EPSPs were triggered by illumination with a blue LED in 2 ms epochs every 3 sec. LED intensity was set to the minimum necessary to evoke the maximum amplitude EPSP. The average baseline EPSP was established with a minimum of 20 sweeps before adding glutamate receptor antagonists (10 μM NBQX (Tocris) and 10 μM CPP (Tocris)) to the bath to block synaptic currents. Responses were monitored as antagonists washed in, and once the EPSP was blocked (∼3 min), a minimum of 20 sweeps were recorded. EPSP amplitudes were calculated by averaging the amplitude 0.5 ms before to 2 ms after the peak of the current.

#### Animal behavior assay

Animals P35 and older were used for the buried cookie assay. Animals were pre-exposed to a reward, specifically an Oreo Mini (Nabisco) chocolate cookie wafer without the icing, 4-5 days before the test was performed. Only animals that ate the cookie within 24 hours during the pre-exposure were used for further testing. Animals were moved to a clean cage and deprived of food overnight to increase motivation. For testing, clean static cages without a water port were filled with approximately 4 cm of fresh bedding and fitted with a clean filter top. During the test, the animals were placed in the clean cage for 10 min to allow them to habituate before returning them to their home cage. Meanwhile, half an Oreo Mini cookie without the filling was buried approximately 2 cm deep in the bedding in a random corner of the test cage. The animal was then placed in the test cage and the timer was started. Up to a maximum of three animals were tested at the same time. A single trial ran for 15 min, or when the animal found the cookie and actively interacted with it by eating it or carrying it away. Animals that failed to locate the reward were shown the cookie and assessed for motivation. They were excluded from the final analysis if they did not interact with the reward. On the conclusion of the trial, the animal was returned to their home cage and provided with food.

#### Transmission Electron Microscopy

Mice were anesthetized with isoflurane and prepared for transcardial perfusion. First, Ringer’s solution (125 mM NaCl, 1.5 mM CaCl_2_, 5 mM KCl, 0.8 mM Na_2_HPO_4_, 20,000 units of heparin (Sigma-Aldrich), pH 7.4) bubbled with 95% O_2_/5% CO_2_ and warmed to 40°C was perfused using a peristaltic pump through the heart, followed by fixative solution containing 4% PFA and 0.25% glutaraldehyde (EMS) in warm PBS. Brains with OBs attached were dissected from the skull and placed in fixative solution at room temperature for 2 hours, then 4°C overnight. The following day, samples were washed three times in fresh PBS, once in PBS with glycine (50 mM) for 10 min, and then returned to PBS. After washing, intact brains were embedded in 4% low-melt agarose and sectioned in the coronal plane at 50 µm. Free-floating olfactory bulb sections were visually screened with fluorescence to identify those containing rat glomeruli.

Olfactory bulb sections containing rat glomeruli were treated with glycine (50 mM, Sigma-Aldrich) and ammonium chloride (50 mM, Sigma-Aldrich) in PBS for 5 min, blocked with 10% albumin for 1 hour, and incubated with primary KsO antibody overnight at 4°C. The following day, sections were washed three times for 20 minutes each in 1% albumin-PBS, incubated with secondary antibody coupled with 1 nm gold particles (Nanoprobe) for 40 min, washed an additional three times in albumin-PBS, and post-fixed with 4% PFA, 2.5% glutaraldehyde, 100 mM cacocylate buffer (Sigma-Aldrich) for 20 min at 4°C. Then samples were washed in the same buffer three times, post-fixed with 1% osmium tetroxide (EMS) and 1.2% potassium ferrocyanide in 100 mM cacodylate buffer for 40 min and washed an additional three times. Sections were dehydrated in an acetone series until 100% (3x) and embedded in Epon resin. Ultrathin sections of 50 nm were obtained with a diamond knife (Diatome) and collected on 300 mesh nickel grids. A silver-enhancement step using Aurion SE-EM kit was performed for 30 min to enlarge the nanogold particles to 8-10 nm size and enhance visualization on the TEM. Finally, sections were post-stained with UranyLess (EMS) for 5 min to improve contrast and observed in a Zeiss Libra 120 operated at 80 kV. High-resolution images (4K x 4K) were obtained with a CCD camera Gatan US4000.

## Data and Code Availability

The datasets and code generated and analyzed during the current study are available from the corresponding author on request.

## Acknowledgements

The authors thank Kathryn Spencer for microscopy assistance, Valentina Lo Sardo and Steffany Dunn for help with iPSC reprogramming and subcloning, and Anastasia Bludova for light-sheet brain imaging. We thank Dr. Qi-long Ying for providing DAC2 and DAC8 rat ESCs. J.W. is a Virginia Murchison Linthicum Scholar in Medical Research and funded by Cancer Prevention & Research Institute of Texas (CPRIT #RR170076) and Hamon Center for Regenerative Science & Medicine. J.C.I.B was supported by the G. Harold and Leila Y. Mathers Charitable Foundation, The Moxie Foundation, The Leona M. and Harry B. Helmsley Charitable Trust (2012-PG-MED002), The Hewitt Foundation, NIH (5 DP1 DK113616) and Universidad Católica San Antonio de Murcia.

## Author contributions

K.K.B., J.W. and J.C.I.B., conceived of the study. K.K.B, J.W. S.K., and P.O supervised the study. K.K.B., J.W., M.K.B.I and B.T.T., designed experiments, wrote, or edited the manuscript. M.S., A.R.R., G.M., and S.K. performed blastocyst microinjection and embryo transfer. R.M.C, P.O., and B.T.T. performed and analyzed the whole-brain imaging experiments. A.L.H., G.L., and B.T.T. performed and analyzed the electrophysiology experiments. K.N.J. and B.T.T. performed and analyzed mouse OB validation histology, imaging, and analysis. M.K.B.I performed immunostaining of chimeric tissues, analysis and quantifications, B.T.T performed all remaining experiments. All authors edited the final drafts.

## Materials and Correspondence

Correspondence and requests for materials should be addressed to K.K. Baldwin and J. Wu.

---

**Supplementary Figure 1: Description of whole-brain volumetric analysis. a,** Schematic of whole-brain imaging and registration workflow. After imaging, brains were aligned to a reference atlas. Subsequently, KsO signal was detected and registered within each defined region. **b,** Representative serial coronal images depict detected KsO signal within registered regions. White outline indicates inset. **c,** Representative flat map projections of KsO signal for both hemispheres of each brain sample, n = 6 animals. **d,** Volumetric analysis and between hemisphere comparison reveals bilateral KsO density asymmetry within animal. Samples registered to the Allen Mouse Brain Reference Atlas. Sample ID (A-F) matches (c).

**Supplemental Table 1: List of Blastocyst Complementation Experiments.** Table includes information on genetic cross used to generate mouse blastocysts, the number of animals generated, and the number identified as chimera based on fluorescent marker (KsO, eYFP) expression. The table also includes the results of injections of rat iPSCs for rescue experiments in mouse with ablate, silence or wildtype (WT) genetic background.

**Supplementary Figure 2: Characterization of development of rat cells in mouse brains. a,** Representative images of cortical CTIP2 staining in rat-mouse chimeras. Insets shown (right) represent the white box, (left). Shown are nuclei (DAPI, blue), rat KsO cells (red) and CTIP2 (green). **b,** The distance to corpus callosum from the nucleus of each CTIP2 positive cell was measured and CTIP2 positive rat cells (red, KsO) were compared to non-red CTIP2 positive mouse cells. Cells from the same section are connected with a black line. The measured distances for mouse and rat cells were averaged per slice. N = 18 slices, 5 animals, and were not significantly different (two tailed, paired t-test). Scale bar 100 μm

**Supplementary Data Figure 3: Description of the optogenetics to study rat-mouse synapses. a,** Expression of pluripotency marker genes Nanog, Oct4, Sox2, and SSEA1 in rat iPSCs carrying channelrhodopsin. Nanog, Oct4 and Sox2 are shown in red and SSEA is shown in green. **b,** Schematic of lentiviral construct inserted in rat iPSCs. Human channelrhodopsin2 fused to the eYFP protein (hChR2H134R0-eYFP) is placed under control of the human Synapsin promoter (hSyn) to enable high expression in neuronal cells. **c,** Traces of light-evoked EPSPs in recorded mouse neurons. Blue triangles mark blue light stimulation. Traces are an average of 20 trials. Scale is 2 mV, 50 ms. n = 10 cells, 7 animals (together with Fig 2f).

**Supplementary Figure 4: Characterization of OSN-compromised mice. a,** Representative images of antibody staining for OMP (green) with nuclei (DAPI, blue) in the OE of E18.5 and P5 WT and Ablate mice. Scale bar, 100 μm. **b,** OMP+ OSNs are depleted in both E18.5 and P5 Ablate OEs. Data shown are mean ± 95% CI, n = 2-3 animals/genotype, 3 slices/animal. Significance was determined by two-way ANOVA followed by Sidak’s multiple comparisons test, \*\*\**p* < 0.001, \*\*\*\**p* < 0.0001. **c,** Representative images showing OMP (green) density in P5 and P120+ mice for both WT and Ablate mice (DAPI, blue). Number of glomeruli are decreased in P5 Ablate mice but not completely absent in adult animals (P120+). **d,** The olfactory bulb (Whole OB) and glomerular layer (GL) areas are smaller in Ablate mice at both P5 and P120+. Data shown are mean ± 95% CI, n = 4-5 animals/genotype, 3 slices/animal. Significance was determined by two-way ANOVA followed by Sidak’s multiple comparisons test, \*\*\*\**p* < 0.0001. **e,** WT mouse PSCs rescue behaviour in both models. Shown are the percentage of the mice that find the cookie, plotted as success percentage against time (s).

**Supplementary Figure 5: Characterization of Rat OSN contribution to olfactory circuits. a,** Rat glomeruli are maintained in the OB in aged mice. Shown are rat KsO (red) and nuclei (DAPI, blue) from a ∼2 year old mouse-rat chimera**. b,** Representative images of transmission electron microscopy of synapses in the olfactory bulb. The electron microscopy images were examined from the area outline in the overview image of the olfactory bulb (left, red is KsO expression from rat cells in a mouse OB). Examples of a mouse-mouse synapses (left) and a rat-mouse synapse (right) are shown.

**Supplementary Figure 6: Characterization of the mouse response evoked by rat neurons. a,** Rat OSN contribution increases cFos expression in the PCx. Shown are the cells positive for cFos per mm^2^ in mouse without any rat contributions. Data shown are the mean ± 95% CI. Significance was tested by One-way ANOVA. \*\*\**p* < 0.0001, \*\*\*\**p* < 0.0003. **b,** Shown, are the cells positive for cFos per mm^2^ for the mice with rat contribution on the contralateral side. Each dot represents data from one hemisphere for both control hemisphere and rat OSN contribution hemisphere within the same animal..**c,** Rat OSN contribution increases Egr1 expression in PCx of Silence animals. Hemispheres with or without rat contribution were stained with anti-Egr1 (white) in serial sections similar to those for which fos staining was performed. **d**, Quantification of cell densities, normalized to the mean density in the no contribution hemisphere for each animal. Data shown are mean ± 95% CI, n = 2 animal/genotype, 15 slices/animal. Significance was tested by Two-way ANOVA followed by Sidak’s multiple comparisons test, \**p* < 0.05.

