## Supplementary Figures for "Building functional circuits in multispecies brains"

### Supplementary Figure 1

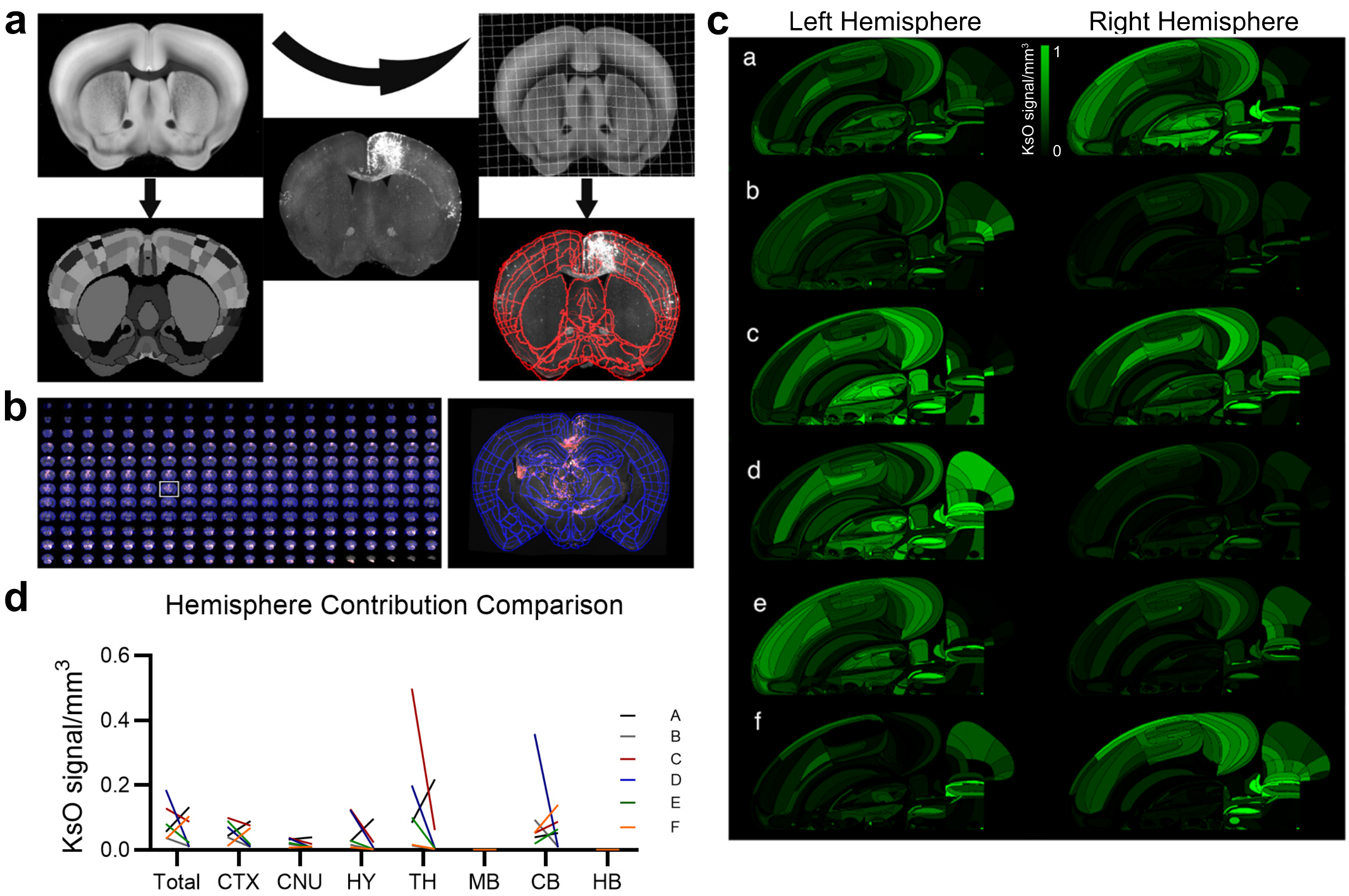

### Supplementary Figure 2

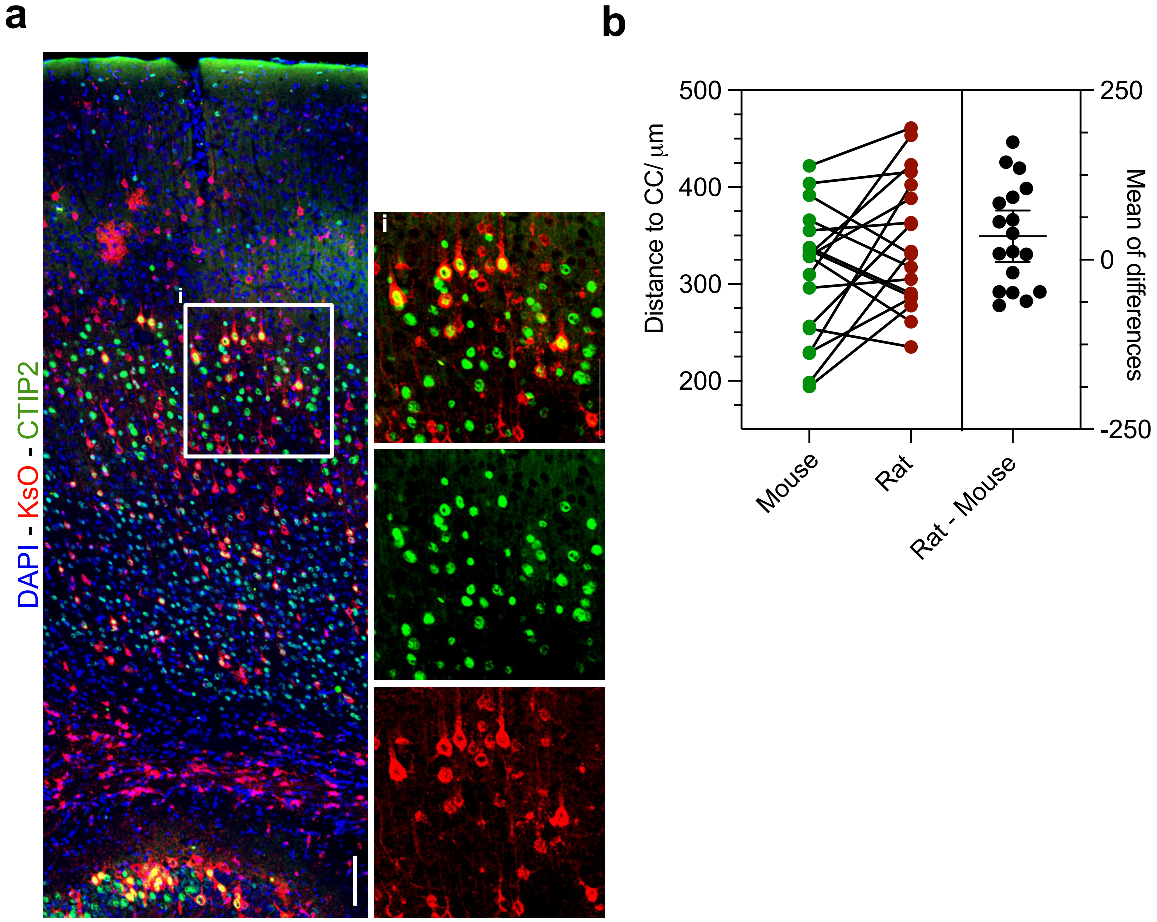

### Supplementary Figure 3

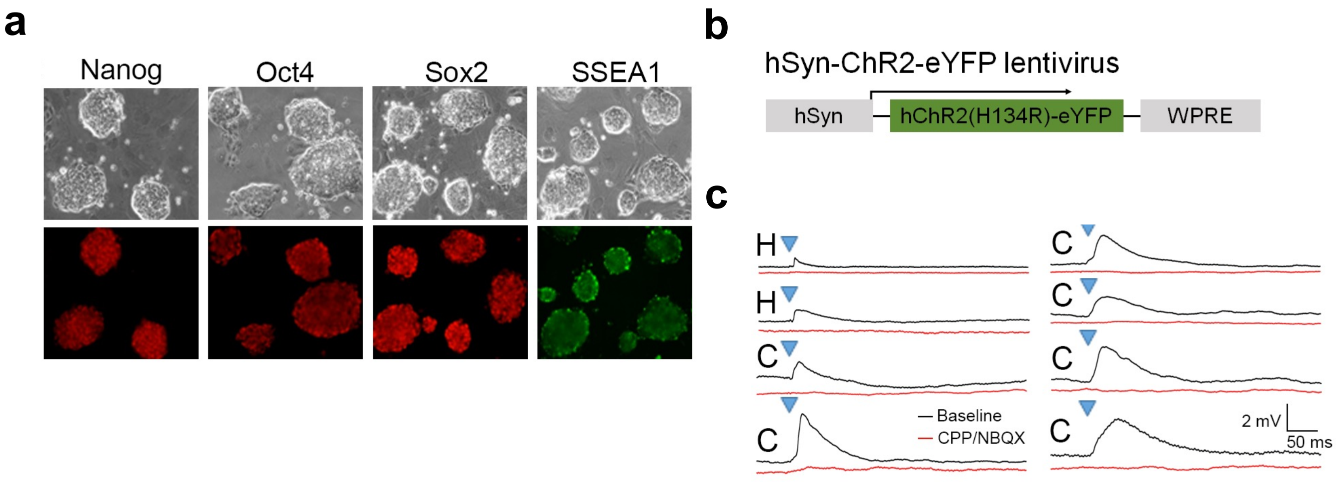

### Supplementary Figure 4

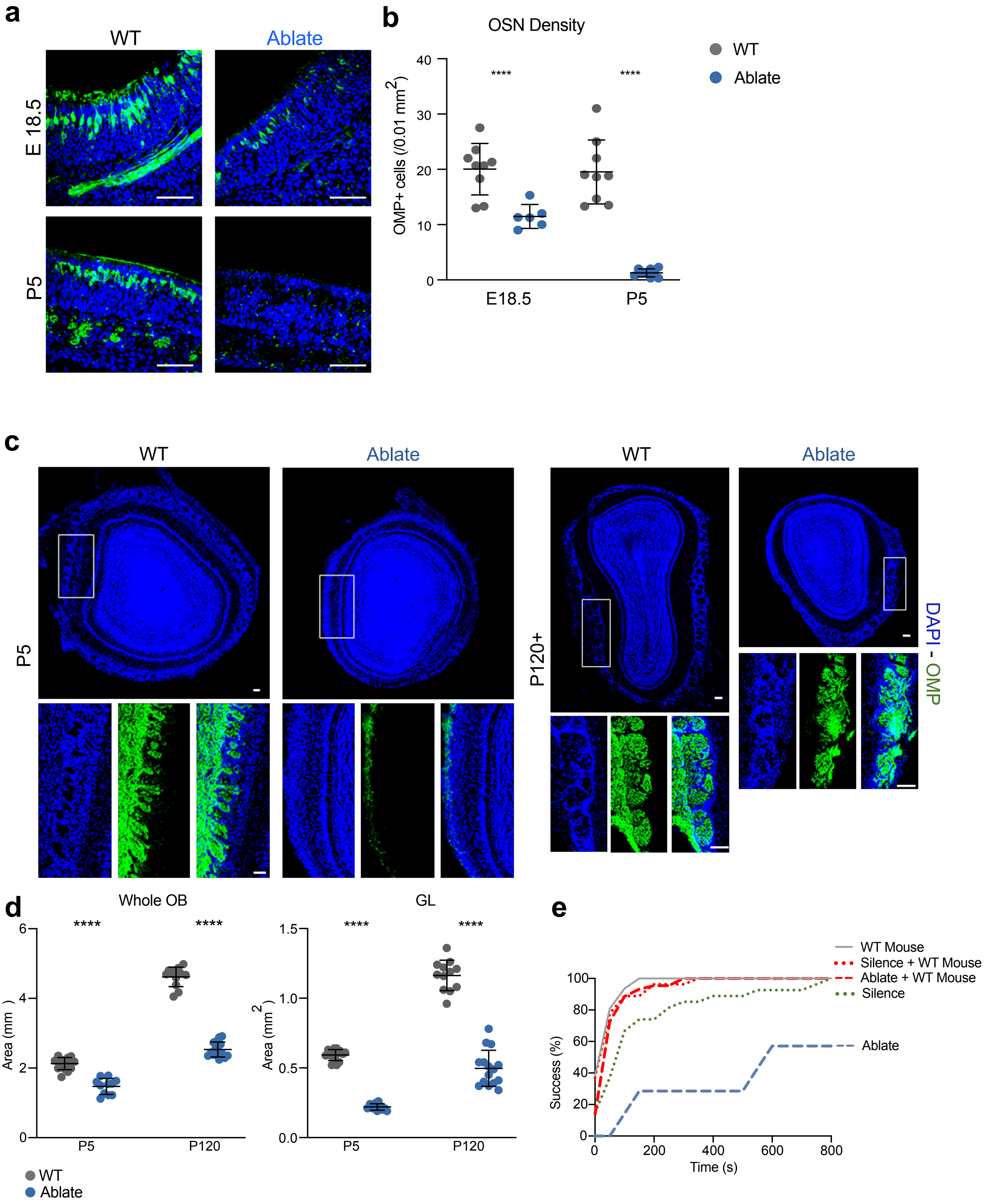

### Supplementary Figure 5

a

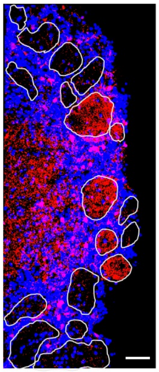

b

Mouse-Mouse Synapse

Rat-Mouse Synapse

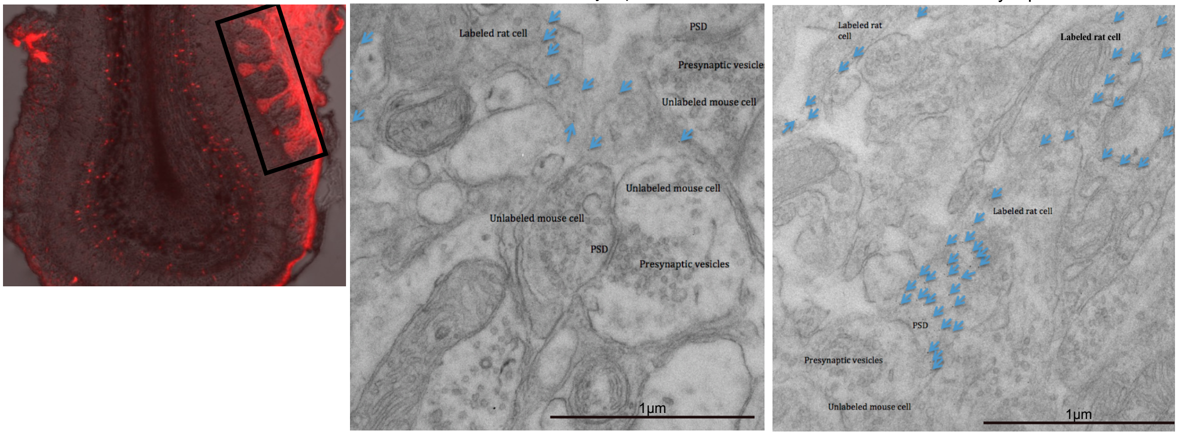

### Supplementary Figure 6

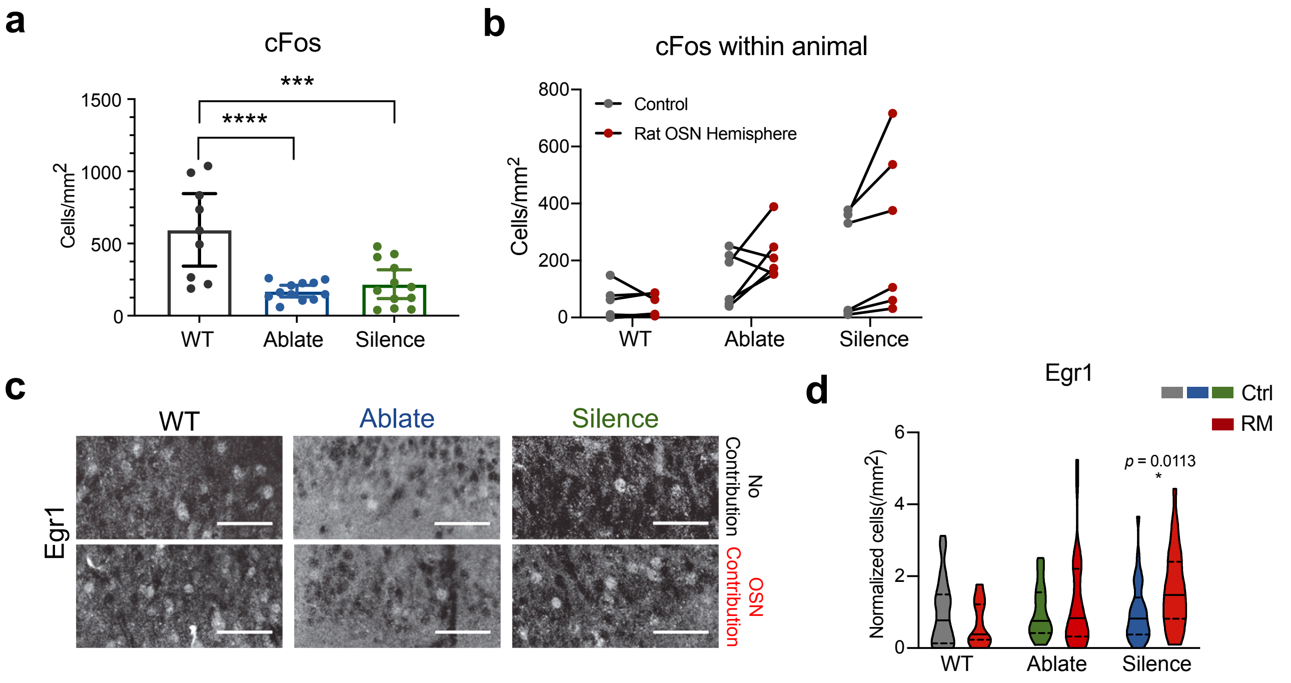
